# Ineffectual Genomic Error Correction Under Environmental Perturbation Dynamically Regulates Mutational Supply and Robustness

**DOI:** 10.64898/2026.03.20.713118

**Authors:** Sashikanta Barik, Parthasarathi Sahu, Koushik Ghosh, Hemachander Subramanian

## Abstract

Adaptive evolution depends on the supply of heritable variation, yet excessive mutation threatens viability by degrading essential molecular functions. Here, we show that this trade-off emerges naturally from the kinetic proofreading mechanism that controls replication fidelity. In our model, environmental shifts alter the optimal driving rate constant of proofreading enzymes, transiently elevating replication error rates and triggering rapid evolutionary change until a new fidelity optimum is reached. This produces alternating periods of stasis and rapid adaptation, consistent with punctuated equilibrium. We further show that coding-region length and population size jointly determine whether adaptation succeeds or mutational collapse occurs, reflecting the balance between mutation supply and error tolerance predicted by the drift-barrier principle. These results provide a molecular mechanistic basis for nearly neutral evolutionary dynamics and illustrate how genomic organization constrains long-term evolutionary resilience.

## 1 Introduction

Evolution proceeds as a complex and nonlinear process shaped by the continual production of heritable variation and the differential preservation of that variation across environments [1]. Because most genetic differences originate from replication errors, the rate at which new variants are introduced—and the extent to which organisms can tolerate their consequences—defines a fundamental axis of evolutionary dynamics [2, 3]. This tension is captured by the interplay between *mutation supply*, which governs the arrival of new variants [4, 5], and *mutational robustness*, which buffers or corrects their potentially deleterious effects. Robustness arises through fidelity mechanisms such as network-level redundancy, and genome features that reduce phenotypic sensitivity to mutation [6, 7]. Too little mutation limits access to adaptive variation, whereas excessive mutation without adequate buffering increases genetic load and extinction risk [8, 9]. Systems with sufficient robustness can accumulate cryptic or nearly neutral variants that become adaptive under environmental change [10, 7], illustrating that evolvability depends not on mutation rate or robustness alone, but on how effectively biological systems maintain a balance between novelty and stability.

In this study, we focus on kinetic proofreading as a tunable, energy-driven mechanism that regulates this balance. Proofreading enhances accuracy by coupling substrate discrimination to non-equilibrium energy consumption, thereby reducing replication or translation errors beyond equilibrium-binding limits [11, 12]. Because proofreading efficiency depends on biochemically adjustable rate constants, it acts as a molecular throttle on mutation supply—allowing cells to modulate fidelity in response to physiological and evolutionary pressures. Under stable conditions, proofreading enforces low mutation rates and supports mutational robustness.

However, this balance is sensitive to the environment. Physical (e.g., radiation, temperature extremes, osmotic or hypoxic stress), chemical (e.g., biocides, heavy metals, antibiotics, reactive oxygen species), and biological stressors (e.g., viral infection, nutrient limitation) all destabilize cellular homeostasis and elevate mutational input [13, 14, 15, 16, 17, 18, 19, 20, 21]. Under such conditions, proofreading functions as the primary intrinsic mechanism that counteracts elevated error rates. Yet proofreading enzymes are themselves encoded in the genome and are therefore vulnerable to stress-induced mutations. As we show here, perturbations can degrade proofreading efficiency at precisely the moment when fidelity is most needed, amplifying mutation supply before selection can re-optimize the underlying biochemical parameters. This dual effect produces a transient burst of elevated mutation followed by a gradual return toward a lower-error state as selection restores proofreading performance.

These molecular disruptions propagate to higher levels of genomic organization. The coding region length determines the mutational target size and sets the baseline mutational burden that proofreading and selection must counteract. Longer coding regions expand the mutational target and increase deleterious load, whereas excessively compact genomes restrict sequence space and limit adaptive potential [22, 23, 8]. Empirical studies show that highly constrained genes tend to be short and compact [24, 25], whereas organisms with relaxed selective pressure tolerate expanded coding and noncoding regions [23]. Thus, an “optimal” coding region length reflects a compromise between maintaining robustness and enabling evolutionary innovation.

Classical theory and comparative analyses clarify how natural selection and genetic drift shape this compromise [22, 26]. Selection refines coding region length and molecular fidelity mechanisms when populations are large enough to distinguish subtle fitness differences, while drift obscures these differences in small populations, allowing mildly deleterious expansions or suboptimal proofreading states to persist or fix [4, 27, 28, 29]. Stress-induced changes in fidelity therefore have different evolutionary consequences depending on population size, determining whether genomes maintain a stable balance between mutation supply and robustness or deviate toward maladaptive trajectories.

To examine the interplay between mutational supply versus robustness, short versus long coding regions, and small versus large population sizes, we develop a theoretical framework that couples replication fidelity to genome evolution using the Hopfield–Ninio kinetic proofreading mechanism [11]. In this framework, the driving-rate constant of the proofreading enzyme determines the replication error rate and admits a well-defined optimum that minimizes mutational input under stable conditions. Environmental perturbations, modeled as temperature shifts, displace the system from this optimum and transiently elevate mutation rates, creating the need for evolutionary re-optimization of fidelity.

By embedding this perturbed proofreading landscape within a population-genetic model, we demonstrate that the error-correcting systems are capable of adapting towards high-fidelity phenotypes even in the absence of guidance from a fitness function. The emergence of such a phenotype through the evolution of coding region length depends critically on population size. Intermediate-length coding regions (defined with respect to mutation rates [30, 31, 32]) exhibit the greatest adaptive capacity, rapidly re-optimizing the driving-rate constant and selection of coding region length; very long regions accumulate excessive mutational burden and tend to collapse under elevated mutation supply; and short coding regions evolve slowly with limited adaptive response, remaining viable only when supported by sufficiently large populations. These results demonstrate that the evolution of coding-region length and proofreading fidelity is not governed by molecular constraints alone but emerges from the interaction between environmental state-dependent mutational supply and the population-size–dependent balance between selection and drift.

## 2 Kinetic Proofreading Model

The error correction mechanism in biochemical processes, like DNA replication, genetic code reading in protein synthesis, and tRNA charging with amino acid, plays a crucial role in preserving and transferring genomic information with high accuracy across generations. The mechanism relies on enzymes discriminating the correct substrates from similar molecules. In the early 1970s, J.J. Hopfield and J. Ninio independently proposed the kinetic proofreading model to explain this discriminatory mechanism [11, 12]. Hopfield’s interpretation of error correction involves the examination of substrate flux (substrate turnover rate) throughout an enzyme-substrate reaction pathway. The enzyme-substrate reaction pathway can be characterized by a series of successive interactions between enzymes and substrates, involving the formation of intermediate complexes, which facilitate the conversion of substrates into products, as illustrated in Fig. (1). Following this reaction pathway, the enzyme effectively incorporates correct substrates while discerning and excluding incorrect substrates to ensure correct product formation.

**Figure 1:**
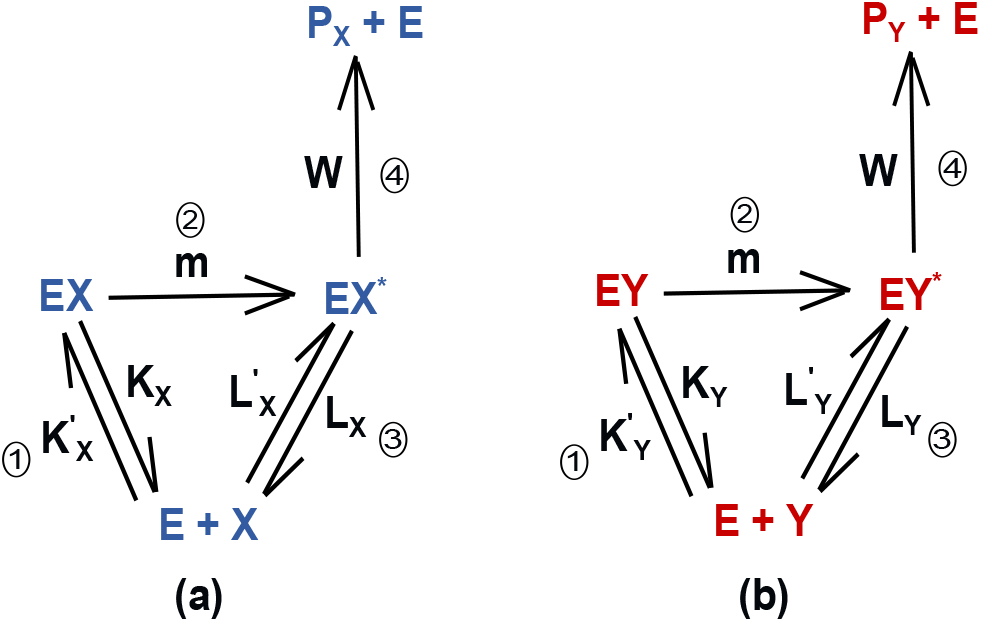
The figure illustrates the correct and incorrect enzyme-substrate reaction pathways with blue colored and red colored chemical participants, respectively. This reaction scheme is identical to the one employed by Hopfield in his seminal kinetic-proofreading paper. E denotes enzyme, while X and Y denote correct and incorrect substrates, leading to formation of P_X_ and P_Y_ as correct and incorrect products, respectively. These pathways encompass multiple reaction steps with intermediate complexes resulting from enzyme and substrate interactions. EX (EY) and EX^*^ (EY ^*^) signify the correct (incorrect) enzyme-substrate complex and intermediate high-energy complex, respectively. 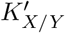 and 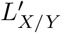 are the association rate constants of intermediate complexes, whereas, K_X/Y_ and L_X/Y_ represent their corresponding dissociation rate constants. Note that the rate constants ‘m’ and ‘w’ are the same for both the reaction pathways. In step 2, ‘m’ signifies the driving rate constant (coupled with external energy source, such as phosphate hydrolysis), while ‘w’ in step 4 facilitates the progression of the intermediate high-energy complex towards product formation before its dissociation through step 3.

Fig. (1) illustrates the enzyme-substrate reaction network, where *X* and *Y* denote the correct and incorrect substrates, respectively. 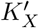 and 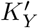 at step 1 represent rate constants for the association of substrates (*X* or *Y*) with the active site of enzymes, which leads to the formation of enzyme-substrate complex (*EX* or *EY*). *K*_*X*_ and *K*_*Y*_ denote the corresponding dissociation rate constants. The driving rate constant ‘*m*’ at step 2, associated with energy consumption via phosphate hydrolysis, drags the enzyme-substrate complex to a high energy transition state. This transition-state complex (*EX*^*\**^ or *EY* ^*\**^) can proceed to product formation through step 4 with rate constant *w* or it can reset the whole reaction by dissociating into free enzymes and substrates through step 3 with a rate constant *L*_*X*_ (or *L*_*Y*_). The association rate constant, 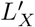 (or 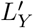), at step 3, is for the thermodynamically uphill, direct transition of enzymes and substrates to high energy intermediate complex (*EX*^*\**^ or *EY* ^*\**^), without energetic assistance.

In Fig. 1, the driving rate constant ‘*m*’ at step 2 and the product-formation rate *w* at step 4 are identical for both pathways and are treated as effectively irreversible transitions, assumptions made by Hopfield as well in his original article [11]. This assumption reflects the biochemical context of DNA replication. During daughter strand construction, DNA polymerase incorporates a dNTP at the growing end of the daughter strand, which forms a phosphodiester bond with the last incorporated nucleotide and releases pyrophosphate (PPi). Rapid hydrolysis of PPi within the polymerase active site or by cellular pyrophosphatases suppresses the reverse reaction, making incorporation effectively irreversible at step 2 [33, 34, 35]. Step 4 is treated analogously: once the product forms, it exits the proofreading network and the system proceeds to the next catalytic cycle, while reversal would require breaking stable bonds and reconstructing the activated intermediate, rendering the backward rate negligible relative to turnover [11, 36].

The difference in the binding strength between the enzyme and the correct and incorrect nucleotides during the initial binding of a dNTP with the polymerase–DNA complex, prior to any conformational change, is relatively small [37, 38, 39]. Furthermore, the enzyme-nucleotide association is largely diffusion-limited, meaning that the rate at which the nucleotide binds is largely determined by how quickly molecules diffuse and collide in solution rather than by strong substrate-specific interactions. Because diffusion governs this step, different substrates (nucleotides) tend to associate with similar rates, and the enzyme cannot strongly discriminate between correct and incorrect substrates during the initial binding event. Consistent with this picture, experiments report only modest (*≈* 2–5-fold) differences in initial binding rates [40, 41]. Larger apparent differences occur only when the “on-rate” reflects a conformational-closing step rather than diffusion-limited binding, in which case, the discrimination no longer arises from proofreading. Therefore, we have assumed the association rate constants to be equal,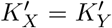 and 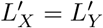 and specificity arises only from the differences in off-rates. Nevertheless, we also examine the effect of moderate variations in association rates on our results in Fig. S10.

Therefore, the discriminatory capability of the reaction arises solely from the difference between kinetic parameters for the *dissociation* of correct and incorrect substrates from enzymes. For effective discrimination, the dissociation rate constants of *EY* and *EY* ^*\**^ should be greater than those of *EX* and *EX*^*\**^, i.e., *K*_*Y*_ > *K*_*X*_ and *L*_*Y*_ > *L*_*X*_. This rate difference impedes the conversion of incorrect substrates into products by trapping them within the enzyme-substrate reaction pathway, while allowing the correct substrates to proceed to product formation.

Within these reaction pathways, steps 1, 2, and 3 in Fig. 1 collectively compose a cyclic path, where step 1 initiates the reaction with the formation of the enzyme-substrate complex, step 2 drives the enzyme-substrate complex unidirectionally to a transition state and step 3 resets the reaction back to enzyme and substrate, which are inputs for state 1, prior to engagement in step 4 for product formation. This cyclic path facilitates the circulation of substrate flux, which is necessary for error correction. The cyclic path entraps the incorrect substrates for a longer time than the correct substrates, resulting in a comparative decrease in the rate of incorrect product formation. Consequently, the greater circulation frequency of incorrect substrates compared to correct ones results in enhanced error correction [42]. This entrapment process is enhanced when the substrates are cycled unidirectionally, enhancing the flux in step 1 - step 2 - step 3 direction, and reducing the flux in the opposite step 3 - step 2 - step 1 direction, which is accomplished by employing an energetically driven stage between steps 2 and 3, which takes *EX* and *EY* to *EX*^*\**^ and *EY* ^*\**^, irreversibly. This thermodynamic drive increases the time delay between the correct and incorrect product formations by increasing the number of times the incorrect substrate cycles through states 1, 2, and 3, compared to the correct substrate, due to their dissociation rate differences.

## 3 Mathematical Model

As discussed in the previous section, the dynamics of substrate flux within the enzyme-substrate reaction pathway plays a crucial role in error correction. To investigate the dynamics of substrate flux, we apply the rate law to the enzyme-substrate reaction pathway, illustrated in Fig. (1), considering both correct and incorrect substrates.

For each substrate *i ∈ {X, Y}*, the enzyme–substrate pathway can be expressed as,

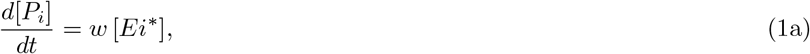

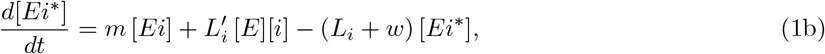

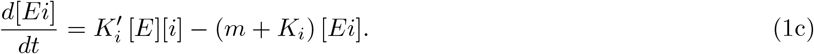

Equation (1a) describes the rate of formation of the correct (and incorrect) product, while Equations (1b) and (1c) describe the formation rates of intermediate complexes for correct (and incorrect) substrates.

We apply the steady-state approximation to the above rate equations for both correct and incorrect enzymatic reaction pathways: 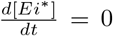 and 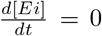. Simplifying the Equations (1b and 1c with the steady-state approximation and defining the error rate (*f*) as the ratio of the rate of incorrect product formation to the rate of correct product formation, we get

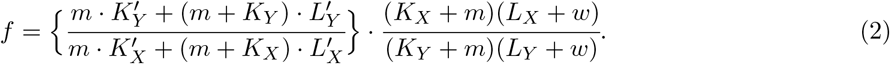

Equation (2) describes the dependence of the error rate on the kinetic parameters of the correct and incorrect reaction pathways. All the kinetic parameters involved in both the correct and incorrect pathways are equal, except the dissociation rate constants.

Furthermore, we have considered the above reaction pathway (Fig. 1) in the absence of the driving step to find out the non-driven error rate.

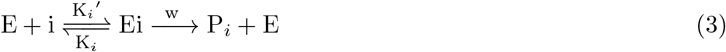

By applying the steady-state approximation to the following rate equations that quantify the dynamics of the above reactions,

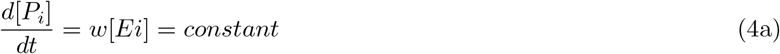

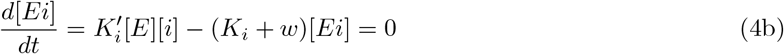

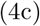

We derive the non-driven error rate as

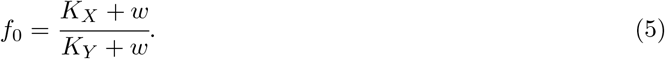

Assuming *w →* 0, the non-driven error rate is,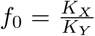. The non-driven error rate shows the energetic discrimination between the correct and incorrect pathways with a single discriminating step.

The error rates (2) and (5) can alternatively be expressed in terms of free energy disparity between the correct and incorrect substrates at the enzyme’s recognition site. The enzyme’s ability to distinguish between correct and incorrect substrates relies on this free energy difference: A greater disparity in free energy enhances the enzyme’s discrimination between correct and incorrect substrates, facilitating error correction and reducing the error rate.

For the reaction pathway with two-step discrimination, step 1 and step 3, illustrated in Fig. (1), the error rate in terms of free energy disparity is given by,

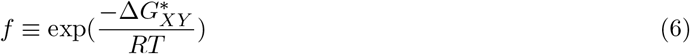

Here, 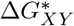 represents the effective free energy difference between correct (X) and incorrect (Y) substrates at the enzyme’s recognition sites throughout the whole reaction pathway [11, 43, 44, 45]. According to Equations (2) and (6), this free energy disparity depends on the rate of dissociation of intermediate complexes in the two discriminating paths. Therefore, an increase in the dissociation rate of the incorrect intermediate complexes amplifies the number of circulations of incorrect substrates in the cyclic pathway formed by steps 1, 2, and 3, leading to an apparent increase in 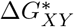. Analogously, the free energy disparity between X and Y at the enzyme’s recognition sites in a single-step discrimination path (Eq. 3) can be denoted as Δ*G*_*XY*_, and the associated error rate *f*_0_ can be alternately written as 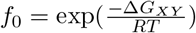.

In Hopfield’s model [11], the total error rate of the enzyme-substrate biochemical pathway with two discriminating paths can attain a minimal value equal to the square of the error rate resulting from the pathway with single-step discrimination 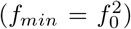, which implies that the maximum value of 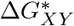 is twice that of Δ*G*_*XY*_. To achieve this minimum error rate and maximum effective free energy disparity, consumption of external energy is required [11, 46, 47, 48], through the driving step 2 of Fig. (1).

Similarly, the external energy consumption can also be quantified in terms of the driving rate constant (*m*). According to the Arrhenius equation [49, 50], ‘*m*’ can be represented in terms of an activation energy barrier (kinetic barrier height *E*_*a*_) between the intermediate complexes.

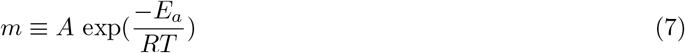

or

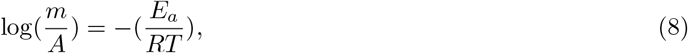

where *E*_*a*_ is the activation energy, A denotes the pre-exponential factor, R is the universal gas constant, and T, the absolute temperature. This energy is required to drag the enzyme-substrate complexes to a high energetic transition state during the driving step 2, which is obtained through phosphate hydrolysis, such as dNTP and GTP hydrolysis, in the case of DNA replication and translation process, respectively [51, 52].

## 4 Evolutionary framework

Our aim here to model phenotypic adaptation of error-correcting enzymes, driven by genetic variation that arises when genomic error correction becomes ineffective under environmental perturbation. We assume that mutations in the perturbed environment primarily affect the catalytic rate *m* of the enzyme, representing the energy-consuming transition step (e.g., phosphate hydrolysis at the active site) that drives proofreading. Although multiple kinetic parameters that characterize the error-correcting enzyme’s function may vary, in reality, many of them do not vary substantially from environmental perturbation to have a noticeable effect on error correction, as we argue below. Moreover, varying a single representative rate captures the dominant adaptive response while avoiding over-parameterization. We therefore treat *m* as the evolving kinetic trait characterizing the enzyme phenotype.

### 4.1 Temperature-shift as environmental perturbation

Environmental perturbations are introduced via a shift in temperature, which predominantly influences the dissociation rate constants in biochemical processes. Higher temperatures destabilize hydrogen bonds between nucleotides, leading to their faster dissociation. The association rate constants 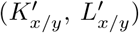 correspond to diffusion-limited encounters between a nucleotide and the polymerase–DNA complex, which therefore produces only modest variation with temperature. In contrast, hydrogen bond dissociation rates vary as *K ∝ e*^*−*Δ*G/RT*^, rendering H-bond lifetimes strongly temperature dependent. In DNA hybridization, for instance, the dissociation rate increases exponentially with temperature, whereas the association rate shows only a weak negative temperature dependence [53]. Thus, association rate constants 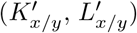 remain comparatively insensitive, whereas dissociation rate constants (*K*_*x/y*_, *L*_*x/y*_) respond strongly to thermal fluctuations.

Although both dissociation rate constants, *K*_*x/y*_ and *L*_*x/y*_, appear in the reaction network, only *K* (*K*_*x*_ and *K*_*y*_) are treated as temperature-dependent in our model. Variations in *L* (*L*_*x*_ and *L*_*y*_) do not affect the error rate as long as their ratio 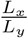 remains fixed. This follows from Eq. 2, where, considering only the step 3 dissociation, the relevant discrimination term appears as 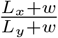; under the assumption *w →* 0, the error rate depends solely on 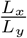. Consistently, the optimal driving constant *m*_0_ is independent of *L* (see Eq. 14).

The temperature dependence of the dissociation rate constant between enzyme and substrate, *K*, follows the transition state theory:

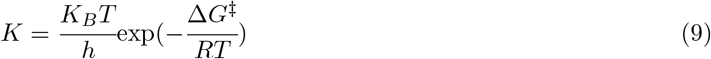

where 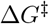 is the free energy of activation for dissociation reaction. Eq. 9 gives *K*_*x*_ = 2*s*^*−*1^ at 293 *K* for the dissociation of correct nucleotides from polymerase during DNA replication [36, 54]. We estimate Δ*G*^*‡*^ = 70.023 *kJmol*^*−*1^, which also aligns with [54].

### 4.2 Genotype-Phenotype Mapping

We next establish a genotype–phenotype relationship to link molecular error rates (genotype) to the evolving driving rate constant (phenotype). Phenotypic change of enzyme is defined as

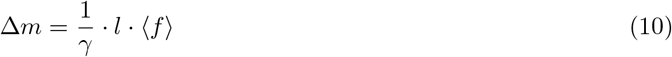

Where, Δ*m* is the change in the enzyme phenotype (which is the change in the driving rate constant in units of *s*^*−*1^, as the result of mutations), *l* is the coding region length, ⟨*f*⟩ is the average mutations per base pair per generation (which is the average error rate of the population in the current generation defined in Eq. 2) and *γ* is a proportionality constant in units of *mutations/generation/sec* (here, *γ* = 1 *mutations/generation/sec*).

Following Eq. 10, 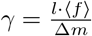, or,

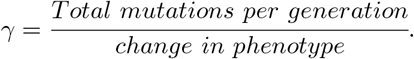

In simpler terms, *γ* quantifies how many total mutations per generation are required to produce a unit change (here, Δ*m* = 1*s*^*−*1^) in the phenotypic driving rate constant.

Equation 10 is motivated by mutation-supply–limited evolutionary dynamics. The quantity *l* ⟨*f*⟩ represents the total number of mutations entering the population per generation (genomic mutation supply). The proportionality constant *γ* converts the number of mutational events into a change in the phenotypic parameter *m*, thereby providing a minimal genotype–phenotype mapping in which evolutionary change of the phenotype slows naturally as mutation supply decreases near the adapted state [55, 56, 57].

### 4.3 Adaptive Dynamics

To observe how mutations shape phenotypic traits in the course of adaptation, we implement the above genotype-to-phenotype mapping to an initial population of *N*_0_ individuals (enzymes), whose trait values, that is, the driving rate constants, (*m*_*individual*_) are normally distributed around the pre-shift optimum (*m*_0,*old*_) with a small standard deviation (*σ* = 0.001) in the initial unperturbed environmental condition (Temperature, *T*_0_). This indicates that individuals have driving rate constants very close to the optimum, thereby maintaining an error rate on the order of 10^*−*8^. A temperature shift Δ*T*, introduced via Eq. 9, increases the replication error rate owing to the absence of immediate phenotypic adjustment. This elevated error rate (Fig. S6) drives phenotypic adaptation toward the new optimum (*m*_0,new_) in the perturbed environment (*T*_new_ = *T* + Δ*T*) through genomic mutations, as described by Eq. 10.

To link error rates to survival, we define the individual elimination probability:

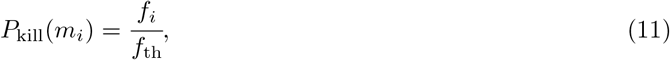

where *f*_*i*_(*m*_*i*_) denotes the individual-specific error rate of the *i*^*th*^ individual, determined by the equation (2) and *f*_th_ (= 10^*−*6^) a threshold error rate beyond which survival is assumed to be improbable. This definition follows from the observation that excessively high mutation rates are typically deleterious [58], i.e., when mutation rates exceed the threshold, essential genes such as those encoding DNA polymerase may accumulate disruptive mutations, effectively silencing or abolishing their activity. This probabilistic mortality function integrates both selection and stochasticity, representing a soft selection filter rather than deterministic elimination, i.e., even maladapted individuals may occasionally survive due to stochasticity. Specifically, high *P*_*kill*_ values correspond to maladaptive regions distant from the low-err optimum, facilitating stronger selective sweeps and faster adaptation rates, whereas low *P*_*kill*_ values correspond to near-optimal phenotypes where drift dominates over selection. Complementing this mortality function, we also define the likelihood of generating new individuals (mutated variants) with trait *m*_*i*_ + Δ*m, m*_*i*_ *−* Δ*m* and *m*_*i*_ into the population; i.e., we define the production probability of viable offspring, *P*_off_, which depends on the relative fitness advantage of surviving individuals. This production probability is therefore defined as a function of selection coefficient *s*(*m*_*i*_), below:

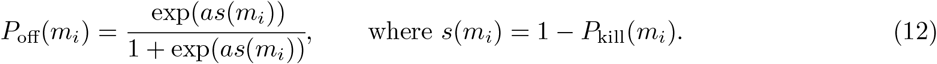

The logistic form was chosen because it provides a bounded, sigmoidal mapping between the selection coefficient and offspring probability, ensuring that individuals with higher survival likelihood (*s →* 1) have a progressively greater probability of producing offspring, while even maladapted individuals (*s →* 0) retain a baseline offspring probability of 0.5, thereby allowing drift to influence evolutionary outcomes. In addition to this baseline behavior, the steepness factor ‘a’ (exponent of Eq. 12) is chosen as 5 (see *Supplementary Information*, Fig. S4) to amplify the discrimination between high- and low-fitness individuals, thereby mimicking the strong selective pressures typically encountered under evolutionary constraints.

Such logistic mappings are widely used in population genetics and quantitative evolutionary models to capture nonlinear selective advantages under finite resources and competition [59, 60]. This offspring probability controls the additions of new mutated variants (*m*_*i*_ + Δ*m, m*_*i*_ *−* Δ*m* and *m*_*i*_) to the population. For instance, if *P*_*off*_ (*m*_*i*_) is high (*≥* 0.75), then two offsprings will be selected at random from the set of three (new variants), ensuring that each of the three possible pairwise combinations occurs with equal probability of 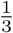. If *P*_*off*_ (*m*_*i*_) is low (*≤* 0.75), then only one offspring will be selected out of the three variants randomly with equal probability of 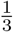.

This offspring probability is stochastic but depend on the selection co-efficient (*s*). However, this choice of the stochastic boundary does not affect our result (see *Supplementary Information*, Fig. S8). This stochasticity is introduced to increase unpredictability and to better match real simulations. Once offspring probabilities are assigned and new individuals are added, the overall population size must also be regulated. If the population exceeds the carrying capacity, *N*_cap_ = 2*N*_0_, individuals are uniformly down-sampled at random, without bias toward higher-fitness individuals. This implements a simple resource constraint, ensuring that the population never exceeds *N*_cap_, which is common in population genetic study.

Following this simulation algorithm, we performed 100 independent simulation replicates to ensure statistical robustness and tracked the evolutionary trajectory of *m*_*avg*_ (*m*_*avg*_ = ⟨*m*_*i*_⟩), population size and adaptation rate across each generation, averaged over all the independent replicates.

## 5 Parameter Regime

To connect molecular kinetics with evolutionary adaptation, we model DNA replication within the proofreading scheme originally proposed by Hopfield. As shown in Fig. 1, *E* represents the DNA polymerase (T7 polymerase), and *X/Y* denote the correct and incorrect nucleotides, respectively. The framework illustrates nucleotide incorporation into the template strand by the polymerase.

In step 1, an incoming deoxynucleotide triphosphate (dNTP) binds to the polymerase at the growing end of the template, forming enzyme–substrate complexes (*EX* and *EY*). In step 2, hydrolysis of the triphosphate releases pyrophosphate, driving the enzyme–substrate complexes into a transition state (*EX*^*\**^ and *EY* ^*\**^). From this transition state, the pathway can proceed either to product formation in step 4 (successful nucleotide incorporation) or to substrate dissociation in step 3. In DNA replication, the baseline error rate (*f*_0_) is typically on the order of 10^*−*4^, whereas the observed minimum error rate (*f*_min_) lies between 10^*−*8^ and 10^*−*9^ [61, 62, 63, 37]. This substantial reduction in error rate is achieved through energy expenditure during step 2.

Here, we have assumed that *K*_*X*_ = *αK*_*Y*_ and *L*_*X*_ = *αL*_*Y*_, *α* represents the energetic bias between correct and incorrect pathways at steps 1 and 3. Furthermore, the association rate constant at step 1 is assumed to be much larger than that at step 3, i.e., 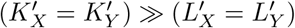, where 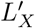 and 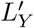 remain small due to the direct transition into the high-energy state (*E* + *X → EX*^*\**^ or *E* + *Y → EY* ^*\**^). Under these constraints, correct nucleotides remain bound to the enzyme’s active site longer than incorrect ones, yielding a higher turnover of correct products. Finally, we set *w* → 0 so that incorrect substrates undergo repeated cycles through steps 1–3, thereby enhancing error correction.

The results presented in this work are robust across a wide range of kinetic parameter values, as long as they satisfy the constraints outlined above. For our analysis, we employed the following representative set of parameters: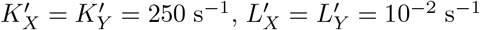, *L*_*X*_ = 0.2 s^*−*1^, and *K*_*X*_ = 2 s^*−*1^ (Table. S1). These values are comparable to the parameters used in previous studies [36, 54], though direct adoption is not possible since those reaction networks differ slightly from ours, which is aligned with Hopfield’s proofreading scheme. Finally, to ensure consistency with the baseline error rate, 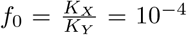, we set the energetic discrimination factor to *α* = 10^*−*4^.

## 6 Results

To examine how environmental perturbations influence the fidelity of genomic processes and the resulting evolutionary dynamics, we first analyze how sudden changes in temperature alter the error-correction efficiency of the proofreading enzyme. Using the Hopfield–Ninio kinetic proofreading framework, we quantify how shifts in temperature destabilize the finely tuned discrimination steps that normally maintain low replication error rates. This disruption elevates the mutation supply, enabling rapid adaptive change until the enzyme evolutionarily re-optimizes its driving rate constant and restores a new error-minimizing configuration under the altered environment. This adaptive mutation scheme leads to the apparent fine-tuning of error rates over long timescales.

Beyond characterizing these fidelity shifts, we investigate how the evolutionary consequences of such perturbations depend on the size of the coding region. Because coding-region length determines both the mutational input available for adaptation and the vulnerability to accumulating harmful changes, evolutionary outcomes reflect a balance between mutational supply and mutational robustness. This balance is mediated by the competition between natural selection—favoring coding region length that improve adaptation—and genetic drift, which erodes weak refinements, particularly in small populations. Together, these forces determine the conditions under which different coding-region lengths can support successful adaptation or collapse under elevated mutational pressure.

### 6.1 Error Minimization by the Optimal Driving Rate Constant

The flux of substrates through the discriminating steps (steps 1 and 3 in Fig. 1) is regulated by the driving step, highlighting the interplay between the driving rate constant and the resulting error rate. At very low *m*, progression from the enzyme–substrate complexes (*EX* and *EY*) to the transition-state complexes (*EX*^*\**^ and *EY* ^*\**^) is comparatively slower than their dissociation. Even the relatively stable *EX* complexes dissociate before advancing, while *EY* dissociates even more rapidly. As a result, both correct and incorrect complexes predominantly revert to free enzyme and nucleotide, leaving step 1 discrimination ineffective and suppressing product formation. Under these conditions, error correction is governed almost entirely by step 3 (see Eq. 13). In contrast, at very high *m*, the reverse problem arises: *EX* and *EY* are rapidly driven into *EX*^*\**^ and *EY* ^*\**^, leaving virtually no complexes at step 1 for differential dissociation to act upon. In this regime, discrimination again relies primarily on step 3, while step 1 contributes negligibly (see Eq. 13).

Thus, at both extremes of *m*, step 1 ceases to contribute to discrimination—either because complexes dissociate before reaching the transition state (low *m*) or because they are driven forward too rapidly for differential dissociation to occur (high *m*). In both cases, error correction is governed solely by step 3, yielding the limiting error rate:

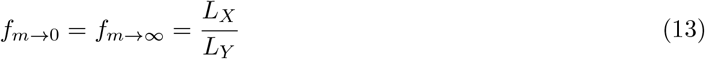

For intermediate values of *m*, however, both steps 1 and 3 are actively engaged in discrimination, lowering the error rate below *α*, the value associated with single-step discrimination. Hence, an intermediate, optimized driving rate constant is crucial for maximizing fidelity. The optimal *m* is obtained by minimizing the error rate (Eq. 2) with respect to the driving rate constant.

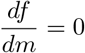

The optimized value for driving constant (*M*_0_) is given by

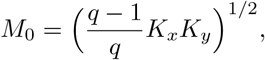

or,

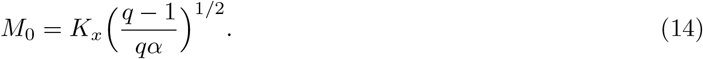

Here,

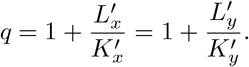

From Equations (2) and (14), the minimum error rate can be approximated, in the limit *w →* 0 and *M*_0_ *≤ K*_*x*_ (see Fig. 2c and supplementary information), as

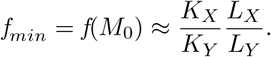

**Figure 2:**
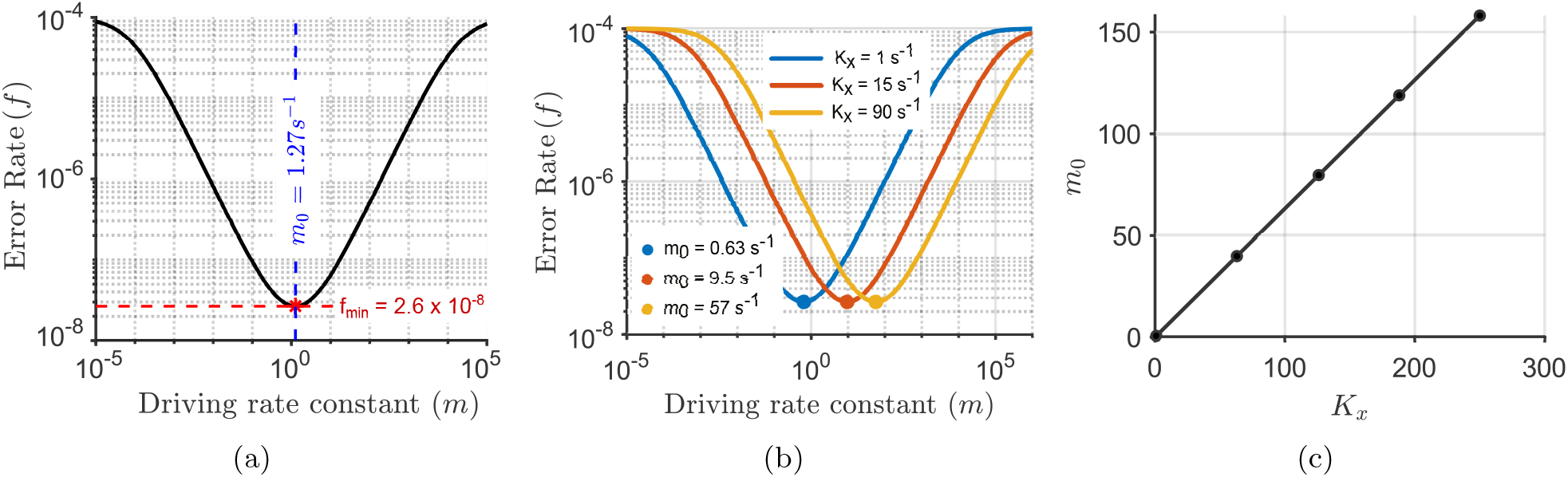
(a) Interplay between the driving rate constant (m) and error rate (f) for K_x_ = 2s^−1^. The red star marks the minimum value of error rate, reflecting the maximum fidelity achieved by the enzyme during proofreading. The associated driving rate constant denotes the optimal energy expenditure for maximal discrimination between correct and incorrect substrates. Saturation at both extremes (low and high m) reflects single-step discrimination (f ≈ 10^−4^), whereas at the optimum (red star), the error rate is reduced to ∼ 10^−8^, consistent with Hopfield’s prediction that the error rate squares. (b) Dependence of error rate on m across different dissociation rates (K_x_). The blue, orange, and yellow curves correspond to K_x_ = 1s^−1^, 15s^−1^, 90s^−1^, respectively. Although the minimum error rate remains constant, the optimal m_0_ shifts to higher values as K_x_ increases, following Eq. 14. Blue, orange, and yellow dots mark the minima at m_0_ = 0.63s^−1^, 9.5s^−1^, 57s^−1^, respectively. (c) Relationship between K_x_ and the optimal driving rate constant m_0_. Increasing K_x_ alters the substrate-flux balance, necessitating higher m_0_ values to restore optimal discrimination.

Therefore, *f*_*min*_ *≈ α*^2^. This result demonstrates that the error rate associated with the reaction pathway involving two discriminating steps is simply the square of the error rate observed in a single-step discrimination pathway. Such a reduction is attainable by investing *optimal* driving energy, provided the conditions of approximation above hold.

The relationship between error rate (*f*) and the driving rate constant (*m*) is shown in Fig.2a. The red star denotes the optimal driving rate constant (*m*_0_), corresponding to the energy investment at which the error rate reaches its minimum. Deviations from this optimum increase *f*, dividing the curve into two sub-optimal regimes. In the left sub-optimal regime (*m* < *m*_0_), *f* decreases as the driving rate constant (*m*) increases, where as in the right sub-optimal regime (*m > m*_0_), *f* grows with further increase in driving rate constant beyond the optimum. At extreme *m* values, *f* saturates at *f*_max_ = *α*, as quantified in Eq. 13. This upper bound (*f ∼* 10^*−*4^) reflects discrimination occurring exclusively through step 3, since step 1 discrimination becomes ineffective. By contrast, at the optimum (*m*_0_, red star), the error rate reaches its minimum (*f*_min_ *∼* 10^*−*8^), where both step 1 and step 3 contribute cooperatively to error correction.

We further observe a systematic shift in the error rate–driving rate constant relationship with varying *K*_*X*_ values, as shown in Fig.2b. While the upper and lower bounds of the error rate remain invariant across *K*_*X*_, the optimal driving rate constant (*m*_0_) increases with *K*_*X*_, consistent with Eq. 14 (see Fig. 2c), while still maintaining the same minimum error rate. This behavior can be understood in terms of substrate flux along the enzymatic pathway. Alterations in *K*_*X*_ disrupt the balance of flux through the two discriminating steps (step 1 and step 3), thereby modulating error rates. Specifically, increasing *K*_*X*_ at a fixed *m* enhances dissociation of [*EX*] and [*EY*], reducing their conversion to [*EX*^*\**^] and [*EY* ^*\**^] and weakening step 1 discrimination. Conversely, lowering *K*_*X*_ diminishes the concentration of [*EX*] and [*EY*], again compromising step 1 discrimination. Thus, to preserve effective discrimination across both steps and ensure prolonged entrapment of incorrect substrates, the driving constant *m* must be tuned accordingly. This requirement highlights the evolutionary optimization of enzymatic energy utilization, enabling replication fidelity to be maintained at the order of 10^*−*8^.

### 6.2 Adaptive Response to Environmental Perturbations

The optimized driving rate constant can be viewed as an evolutionary trait shaped by the kinetic parameters of genomic error correction processes. These kinetic parameters are not fixed but vary with environmental conditions. For instance, temperature fluctuations alter dissociation rate constants in fig. 1, with higher temperatures generally accelerating the dissociation. In our framework, we model temperature variation as the environmental perturbation by adjusting the dissociation rate constants, particularly those of step 1 (*K*_*X/Y*_), within the biochemical pathway. Because dissociation rates are environment-dependent (see Methods), the optimal driving rate constant required to minimize error is itself dynamic. To maintain a minimal error rate (on the order of 10^*−*8^ in DNA replication), enzymes must therefore recalibrate their driving rate constant under changing environment. This selective pressure drives the evolutionary fine-tuning of the trait, ensuring that the driving rate constant remains optimally adapted to the prevailing environment.

To examine this process, we implement a mechanistic adaptive model *without imposing a predefined fitness function* (see *Evolutionary Framework section*). Starting from an initial population of 10^4^ individuals, we expose the system to an abrupt temperature shift from *T*_old_ = 293K to *T*_new_ = 335K. This perturbation is incorporated by altering the dissociation constant of step 1, with *K*_*x*_ increasing from *K*_*x*,old_ = 1s^*−*1^ to *K*_*x*,new_ = 42s^*−*1^, according to Eq. 9. The sudden change raises the population’s mean error rate by roughly ten-fold relative to its initial minimum value, as the enzyme cannot immediately readjust its driving rate to the newly required optimum (*m*_0,old_ = 0.632s^*−*1^ versus *m*_0,new_ = 26.569s^*−*1^). This elevated replication error, in turn, generates genetic variation across the coding region of the genome (assumed length 10^6^ bp), which then manifests as phenotypic changes in the enzyme that may prove either beneficial or deleterious.

Mutations, arising as the immediate consequence of the elevated replication error, generate the genetic variation that drives adaptation. Following this adaptive phase, the enzyme progressively reduces the elevated error rate and restore functionality under the perturbed environment. The pace of this adaptive response is governed by the mutation rate, defined here as the average number of mutations per base pair per generation. Higher mutation rates accelerate adaptation by providing more variants for selection, whereas lower mutation rates slow the process. As the population approaches the minimum error, the effective mutational input diminishes, and enzymes are progressively selected to operate at the newly required optimal driving rate.

This enzymatic evolution, summarized in Fig. 3a, highlights how error rate shifts under environmental perturbation. In the stable environment (*K*_*x*,old_ = 1 s^*−*1^), the error correction dynamics is illustrated by the blue curve with the blue dot marking the maintenance of high fidelity at the optimal driving rate constant (*m*_0,old_ = 0.632 s^*−*1^). After an environmental shift, this fidelity is disrupted: the enzyme, still operating with the old driving rate, incurs a higher error rate, as represented by the blue dot displaced onto the orange curve corresponding to the perturbed environment (*K*_*x*,new_ = 42 s^*−*1^). Because the enzyme cannot instantly readjust its driving rate to the new optimum, replication becomes error-prone, introducing genetic variation that fuels evolutionary change. Over successive generations, adaptation drives the enzyme toward the new optimum (*m*_0,new_ = 26.569 s^*−*1^), depicted in the third panel by the transition from the blue to the orange dot on the orange curve, thereby restoring high fidelity under the new conditions.

**Figure 3:**
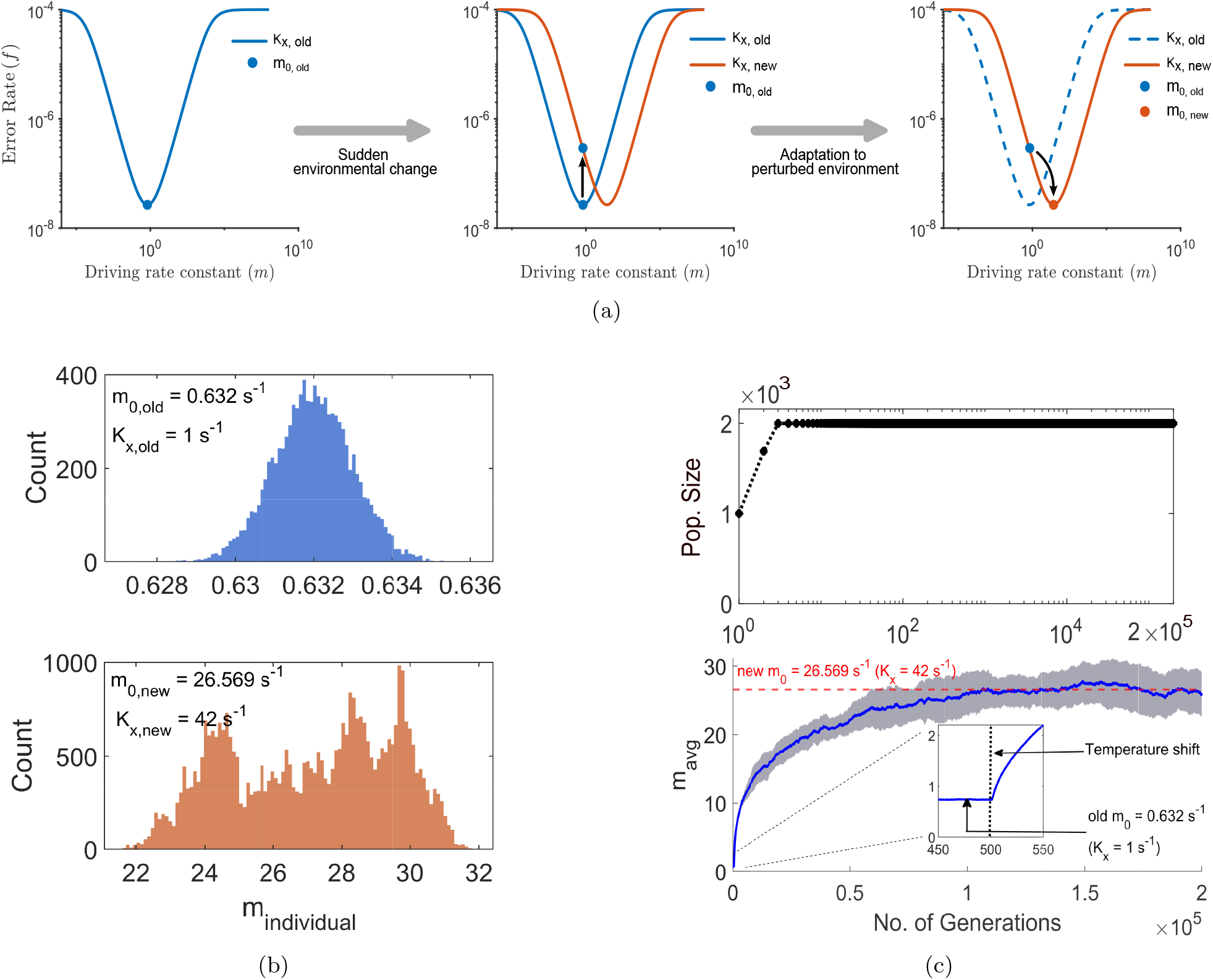
Enzymatic adaptive dynamics to minimize error rate in a perturbed environment. Achieving this minimum error rate requires enzymes to operate at a driving rate constant tuned to the kinetic parameters dictated by prevailing conditions. (a) Schematic representation of enzymatic adaptation due to an abrupt perturbation. Initially (first panel), the blue dot on the blue curve marks the optimal driving rate constant, corresponding to a minimum error rate (f_min_ ≈ 10^−8^) in the unperturbed environment (K_x,old_ = 1 s^−1^, T_old_ = 293 K, m_0,old_ = 0.632 s^−1^). After the environmental shift, the error rate rises about ten-fold because the enzyme cannot immediately adjust to the new optimum. This is shown in the second panel, where the blue dot shifts from the blue curve to the orange curve (error rate vs. driving rate constant). Where the orange curve represents the perturbed environment (K_x,new_ = 42 s^−1^, T_new_ = 335 K, m_0,new_ = 26.569 s^−1^). Over successive generations, mutations drive adaptive evolution toward the new optimum. The third panel depicts this transition, as the blue dot evolves into the orange dot on the orange curve, representing the adapted state. (b) Distribution of individual optimal driving rates (m_individual_). The blue histogram represents the initial population of 10^4^ individuals with mean m_0,old_ = 0.632 s^−1^. The orange histogram shows 4 × 10^4^ individuals after 2 × 10^5^ generations, with mean m_0,new_ = 26.569 s^−1^, indicating successful adaptation. (c) Evolutionary trajectory of adaptation. The blue curve (with 99.9% confidence interval) shows the mean driving rate constant over generations. The trajectory saturates at the new optimum (red horizontal line) without an explicit fitness function, reaching steady state with minimal error rate in ∼10^5^ generations. The corresponding population size dynamics are shown in black (with population axis in logarithmic scale for visualization of the early rise).

This adaptive process is further quantified in Fig. 3c and Fig. 3b, based on Monte Carlo simulations of trait evolution under mutational bias, performed *without an explicit fitness function*. Fig. 3b shows the distribution of individual driving rates (*m*_individual_) within the population: the blue histogram corresponds to the initial population in the unperturbed environment (*N* = 10^4^, mean *m*_0,old_), while the orange histogram shows the adapted population in the perturbed environment (*N* = 4 × 10^4^, mean *m*_0,new_). The evolutionary trajectory of this transition is captured in Fig. 3c, where the blue curve depicts the progression of the mean driving rate (*m*_avg_) across generations. The trajectory converges to the new optimum (red horizontal line), attaining a steady state with minimal error rate after approximately 10^5^ generations, while the corresponding population size dynamics are shown in black.

To quantitatively characterize this trajectory, we define the adaptation rate as the change in phenotype per generation, which is computed directly from simulations as the per-generation change in *m*_avg_ of the population. As shown in Fig. 4, the adaptation rate shows an initial sharp rise reflecting the sudden error increase imposed by the temperature shift, followed by a gradual decline as mutation rate decreases and the population mean converges toward the new optimum (*m*_0,new_). This observation is in agreement with the core assumptions of Fisher’s geometric model, in which Δ*m* is taken to be uniform across the population within a generation and the mutational step size diminishes as the phenotype approaches its optimum. A comparison of evolutionary dynamics under uniform Δ*m* versus individual-specific Δ*m* is provided in Fig. S1 of the Supplementary Information.

**Figure 4:**
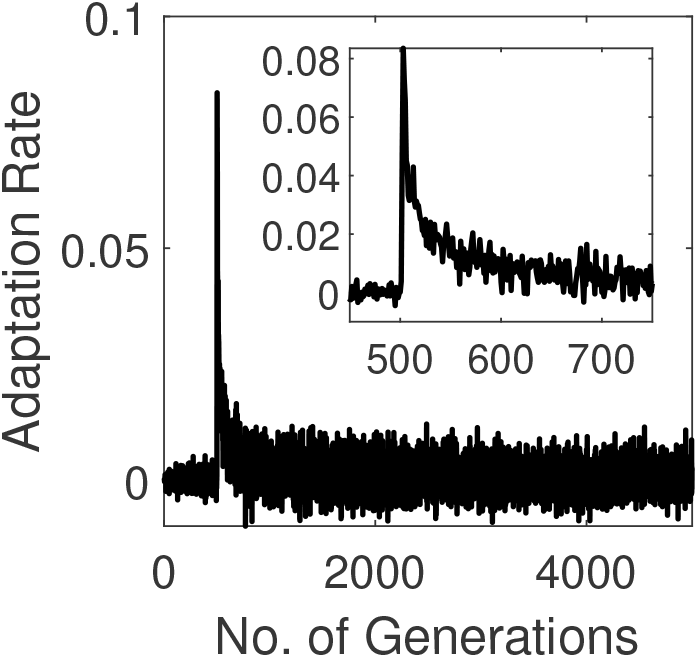
Evolution of the adaptation rate, defined as the change in phenotype m per generation, across generations. The adaptation rate rises sharply in the early generations just after the perturbation at 500^th^ generation, reflecting the sudden increase in replication error following the environmental (temperature) shift. As the population mean trait approaches the new optimum, the adaptation rate progressively declines, driven by the reduced mutation rate at steady state.

A temperature shift increases error frequencies in replication and protein synthesis, and errors generated in one process propagate into others, elevating organism-level mutation supply [64]. The resulting surge in mutational input produces rapid phenotypic change, corresponding to the high initial adaptation rate observed in Fig. 4. As enzymes readjust their driving rate constant and restore fidelity, mutation supply declines and the population mean approaches the new optimum *m*_0,new_, causing the adaptation rate to slow and the system to enter evolutionary stasis. Subsequent environmental disturbances repeat this cycle of rapid change followed by stabilization, giving rise to alternating phases of evolution and stasis consistent with punctuated-equilibrium–like dynamics [65]. This signatures of punctuated equilibrium is illustrated in Fig. S13.

### 6.3 Influence of Coding-Region Length and Population Size on Adaptive Dynamics

To examine how genome size influences evolutionary dynamics under changing environments, we investigated the fate of populations differing only in the length of their coding region. Using the same evolutionary framework described earlier, we simulated populations with coding-region lengths of 10^3^, 10^6^, and 10^8^ bp, each exposed to environmental perturbations that alter the optimal driving rate constant of the proofreading enzyme (Fig. 5). The perturbation was introduced at the 500^th^ generation by shifting the environmental temperature, which increases the replication error rate by displacing the error-correcting enzyme from its optimal configuration.

**Figure 5:**
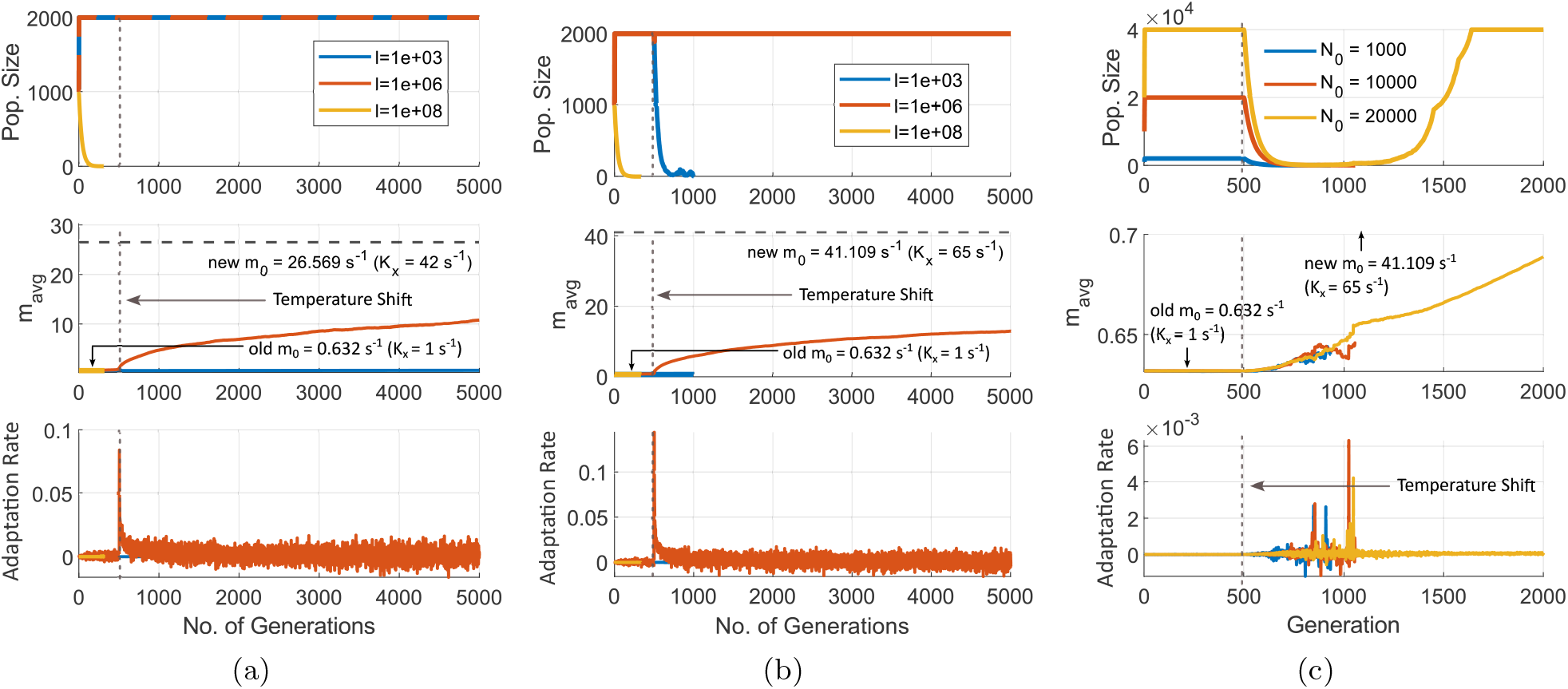
The figure illustrates the evolutionary dynamics of populations with varying coding region lengths and initial sizes under environmental perturbations. Each panel shows population size, mean driving rate constant (m_avg_) of the error-correcting enzymes, and adaptation rate. **(a)** Populations with initial size N_0_ = 10^3^ and coding-region lengths of 10^3^, 10^6^, and 10^8^ bp (blue, orange, and yellow curves, respectively) subjected to a temperature shift of ΔT = 42 K at generation 500. Populations with very long coding regions (10^8^ bp) collapse prior to perturbation due to rapid accumulation of deleterious mutations. Intermediate-length populations (10^6^ bp) retain sufficient mutational input to re-optimize efficiently after perturbation. Short coding-region populations (10^3^ bp) survive the perturbation but adapt slowly because of limited variation. **(b)** The same populations exposed to a stronger perturbation of ΔT = 47.68 K. Under this harsher shift, short coding-region populations (10^3^ bp) collapse, as the elevated error rate overwhelms their restricted adaptive potential. Intermediate-length populations (10^6^ bp) continue to re-optimize, while very long coding regions (10^8^ bp) collapse even before perturbation. **(c)** Evolutionary dynamics of populations with a fixed coding-region length of 10^3^ bp but different initial population sizes (N_0_ = 10^3^, 10^4^, and 2 × 10^4^; blue, orange, and yellow curves) under the strong perturbation (ΔT = 47.68 K). Larger initial populations provide sufficient mutational input and buffer against drift, enabling recovery and re-optimization. When N_0_ = 2 × 10^4^, the population successfully adapts despite its small coding-region length, whereas smaller populations collapse.

Our results reveal a clear selection for specific coding lengths, disfavoring smaller and larger lengths and selecting an intermediate length, depending on the mutational rate and population size. Populations with very long coding regions (10^8^ bp) accumulate deleterious mutations so rapidly that they collapse even before any perturbation occurs (*Muller’s ratchet*), illustrated in Fig. 5a and Fig. 5b. The large mutational target associated with longer genomes dramatically elevates the probability of disruptive substitutions. This behavior illustrates that an excessive mutational supply does not necessarily enhance evolvability; in small or moderately sized populations, it instead accelerates mutational meltdown.

In contrast, populations with intermediate coding-region length (10^6^ bp) maintain low error rates under stable conditions and exhibit rapid adaptation when perturbed (Fig. 5a). After the environmental shift, the average driving rate constant *m*_avg_ increases quickly toward the new optimum, and the adaptation-rate curve shows a sharp transient rise, indicating efficient acquisition of beneficial mutations. This coding-region length provides a sufficient mutational supply to explore adaptive variants while keeping the accumulation of harmful changes at a manageable level, thereby enabling robust evolutionary response.

Populations with the shortest coding region (10^3^ bp) display a different pattern. Under the perturbation of temperature shift of 42 K, these populations remain viable but evolve extremely slowly: their small coding region produces very limited mutational variation, resulting in an almost steady trajectory of the average driving rate constant with negligible phenotypic change, shown in Fig. 5a. When the perturbation is stronger (Δ*T* = 47.68 K) in Fig. 5b, the population experiences a sharp increase in the replication error rate. This elevated mutation rate, coupled with limited adaptive potential arising from the small coding region, results in negligible phenotypic change, as reflected in the evolutionary trajectory of *m*_*avg*_ and adaptation rate plot of Fig. 5b. Consequently, the population undergoes extinction, unable to accumulate sufficient beneficial mutations for survival under the new environmental conditions.

Nevertheless, this short coding region becomes selectively favored in large populations, where a subset of individuals survives by maintaining error rates well below the threshold (*f*_*th*_), effectively lowering their elimination probability (*P*_*kill*_) and persisting long enough to accumulate beneficial mutations. These survivors gradually move away from the critical error boundary and adapt slowly toward the new optimal driving rate (Fig. 5c). The small coding region becomes selectively favored not because it inherently promotes adaptation, but because a sufficiently large population compensates for its limited mutational input. This demonstrates that slow, nearly neutral adaptation can be deleterious and lead to extinction in small populations, whereas the same trait can be naturally selected in large populations, where selection effectively amplifies even subtle fitness advantages. Thus, both coding region length and population size jointly determine evolutionary resilience and the likelihood of extinction under environmental stress.

Together, these results (including Fig. S7 and S9) demonstrate that the evolutionarily favored coding-region length is not fixed but emerges from the balance between mutation supply and mutational robustness, as modulated by population size. Very long coding regions fail because they overload the population with deleterious mutations; intermediate coding regions support rapid and stable adaptation; and very short coding regions survive only when large population sizes provide enough mutational input to enable evolutionary response. This interplay between coding region length, mutation rate, and population size directly reflects the competition between natural selection and genetic drift that shapes molecular evolution.

## 7 Discussion

In this study, we show how living systems dynamically regulate mutational supply, needed for adaptation, with robustness to mutations, to avoid error catastrophe, without foresight, that is, without the help of an explicit fitness function. Error-correcting systems maintain robustness by lowering the mutational input. However, environmental perturbations disturb this balance by increasing the mutational supply through destabilization of the error-correcting systems, which helps accelerate adaptation to the said environmental perturbation. Once adapted to the new environment, the error-correcting system (polymerase) with altered driving rates again achieve low error rates, thereby regaining robustness.

Our model provides a mechanistic framework for this dynamic regulation of mutational supply by explicitly linking replication fidelity to environmental perturbation through kinetic proofreading. Replication fidelity arises from an intrinsic trade-off between the driving rate constant and the resulting error rate, as illustrated in Fig. 2. Similar trade-offs between error rate and speed or energy cost are observed in Ref. [36, 67]. Environmental perturbations—such as temperature shifts—modify the kinetic parameters that govern substrate binding, dissociation, and discrimination, thereby displacing the enzyme from its optimal operating point, where the driving rate constant is tuned to minimize errors. Once perturbed, the mismatch between correct and incorrect substrate flux destabilizes the proofreading cycle, causing an abrupt rise in replication errors and increasing the mutational input to the population. This transient surge in mutation supply generates abundant genetic variation, enabling rapid adaptive change under the new environmental regime before fidelity is re-established. As the enzyme evolves toward a new driving-rate constant that restores a low error rate, the mutation rate declines, and the system settles into a renewed phase of evolutionary stasis. This sequence of perturbation, accelerated adaptation, and subsequent stabilization quantitatively mirrors the dynamics of punctuated equilibrium observed across microbial, molecular, and phenotypic evolution [68, 69, 70, 71, 72].

The above-mentioned balancing of mutational supply and robustness depends on coding sequence length and population size. Extending the coding region enlarges the mutational search space, increasing the probability of generating beneficial changes while simultaneously increasing exposure to harmful substitutions within essential sequence positions. Finite populations can negotiate this trade-off: they selectively retain intermediate coding lengths that allow rapid adaptation without incurring catastrophic error accumulation. Our results illustrate this dynamic clearly: populations with very long coding regions accumulate detrimental mutations so rapidly that they collapse even before any environmental perturbation occurs. Populations with intermediate-length coding regions maintain low error rates under stable conditions and adapt efficiently following a perturbation. In contrast, populations with very short coding regions exhibit extremely slow, nearly neutral evolutionary change because their limited mutational input restricts adaptive potential. When the perturbation is strong and the induced mutation rate becomes high, these small-coding-region populations collapse unless they are supported by sufficiently large population sizes, which provide the increased mutational supply necessary for adaptation (see Fig. 5). This scaling behavior parallels the drift-barrier hypothesis, which states that the optimization of molecular traits is limited by the point at which further refinements are too subtle to overcome the stochastic noise imposed by drift [8, 66].

The same principles may help explain observed cases of rapid evolutionary change, such as the emergence of antibiotic resistance, adaptive melanism, and phenotypic shifts in invasive or climate-stressed species. Because environmental perturbations directly influence molecular fidelity mechanisms, they can modulate evolutionary rates across ecological and geological timescales alike. Thus, this framework provides not only a mechanistic understanding of how selection, drift, and molecular fidelity interact but also a predictive foundation for assessing the adaptive capacity and long-term resilience of evolving populations under fluctuating environments.

## Author Contributions

HS conceived the idea. SB, PS, and KG, together with HS, researched and wrote the article.

## Data Availability

All data necessary to reproduce the results are contained in the manuscript, and the associated MATLAB codes are available at: https://github.com/Sashik05/kinetic-proofreading-mutation-robustness.

## 1 Optimized driving rate constant

Our study establishes a relationship between the error correction efficiency of an enzyme (polymerase) and environmental stress. We quantify the ineffectiveness of the error correction process due to sudden environmental perturbations, and the eventual adaptation of the enzyme to the new environment. To regain high accuracy after a perturbation has occurred, enzymes evolve to adapt the newly required optimized driving rate constant, *M*_0_, in the perturbed environment.

The optimized driving rate constant *M*_0_ can be obtained by minimizing the error fraction, equation (3), w.r.t the driving rate constant, using

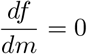

Assuming 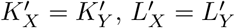 and *w* = 0, we get

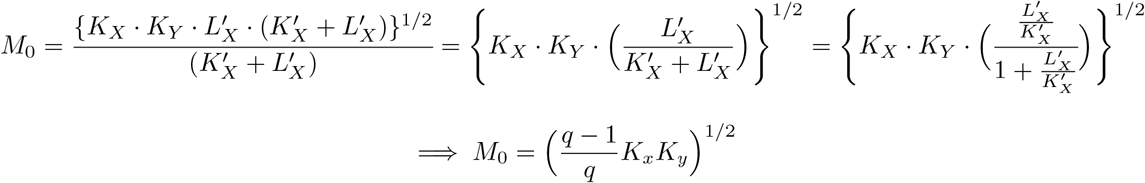

Or,

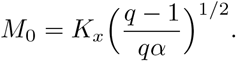

where,

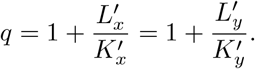

## 2 Calculation of minimum error fraction

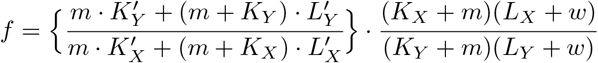

Applying the approximations *W →* 0 and 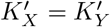,

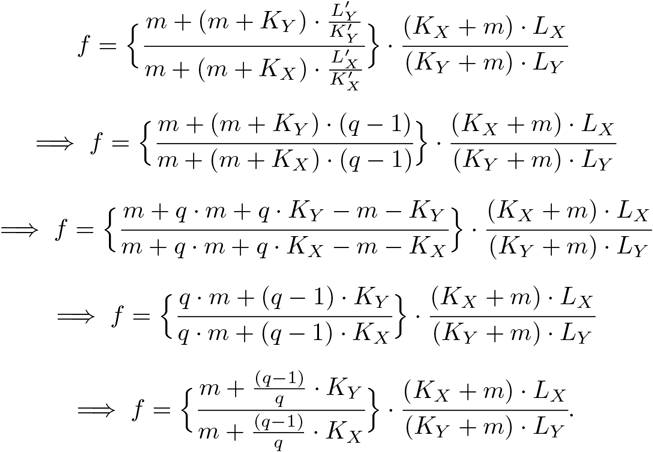

Implementing the optimized driving rate equation, equation (17), in the above equation,

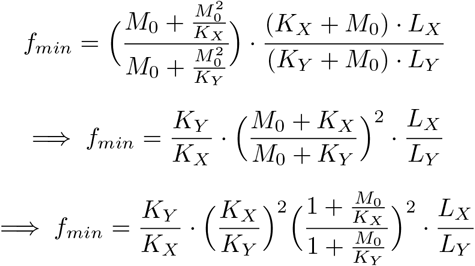

*M*_0_ is the optimized driving rate constant that balances the discriminatory substrate flux between the two discriminating paths, step 1 and step 3, resulting in error minimization. This indicates the driving rate constant *M*_0_ should be less than the *K*_*X/Y*_ for effective discrimination through step 1. Otherwise, [*EY* ^*\**^] will increase, promoting an increase in the rate of incorrect product formation, which increases the error fraction. Therefore, we assume *M*_0_ ≲ *K*_*X*_ and *M*_0_ < *K*_*Y*_. This leads to

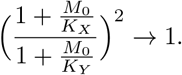

Now, *f*_*min*_ will be,

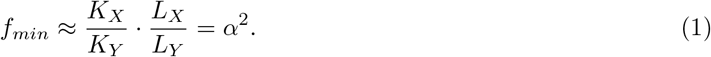

### Algorithm 1: Simulation of trait evolution of the population with fixed coding region length under kinetic proofreading–based adaptation dynamics

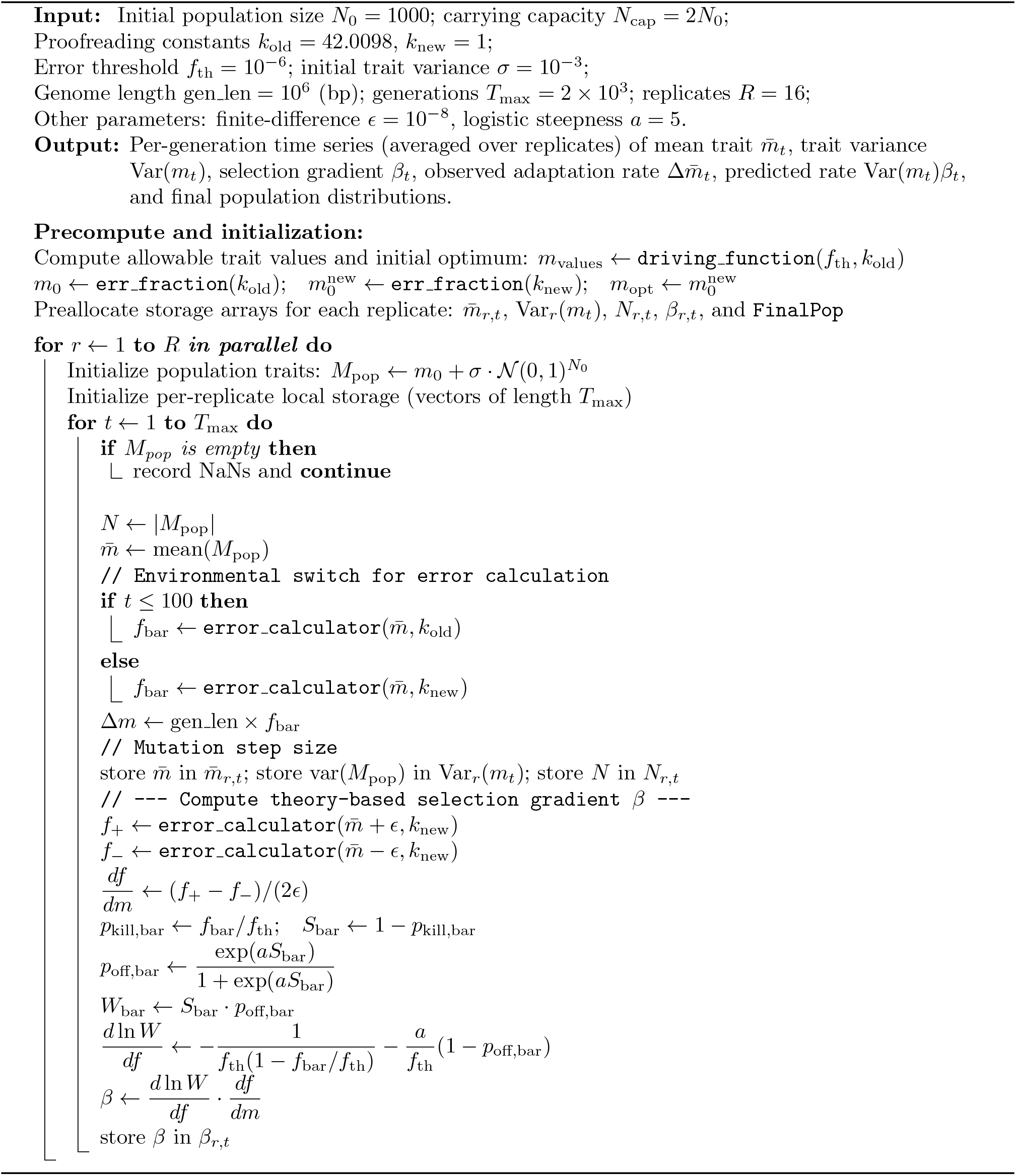

### Algorithm 2: Birth–death and mutation dynamics per generation (Cont.)

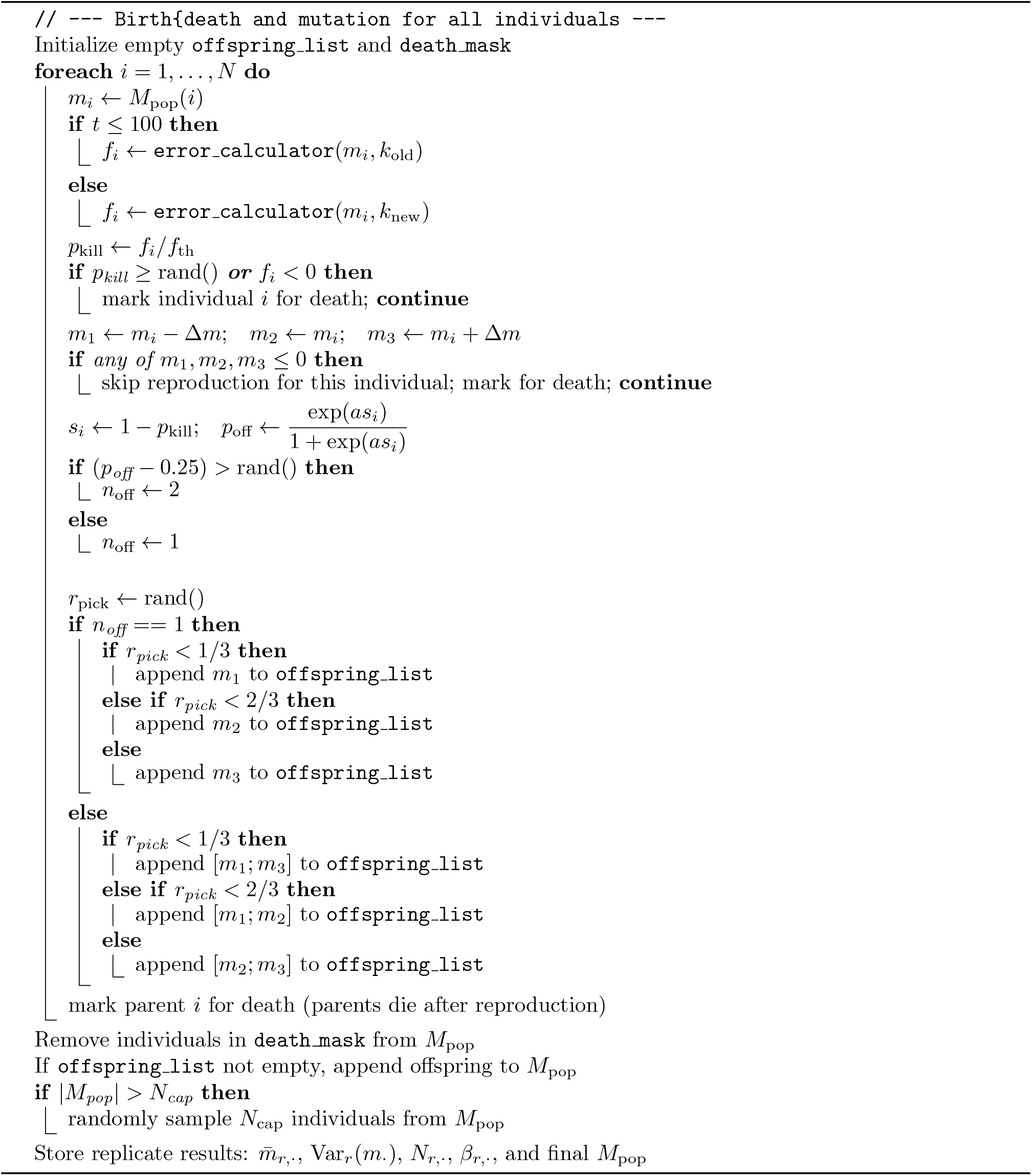

### Algorithm 3: Postprocessing and plotting

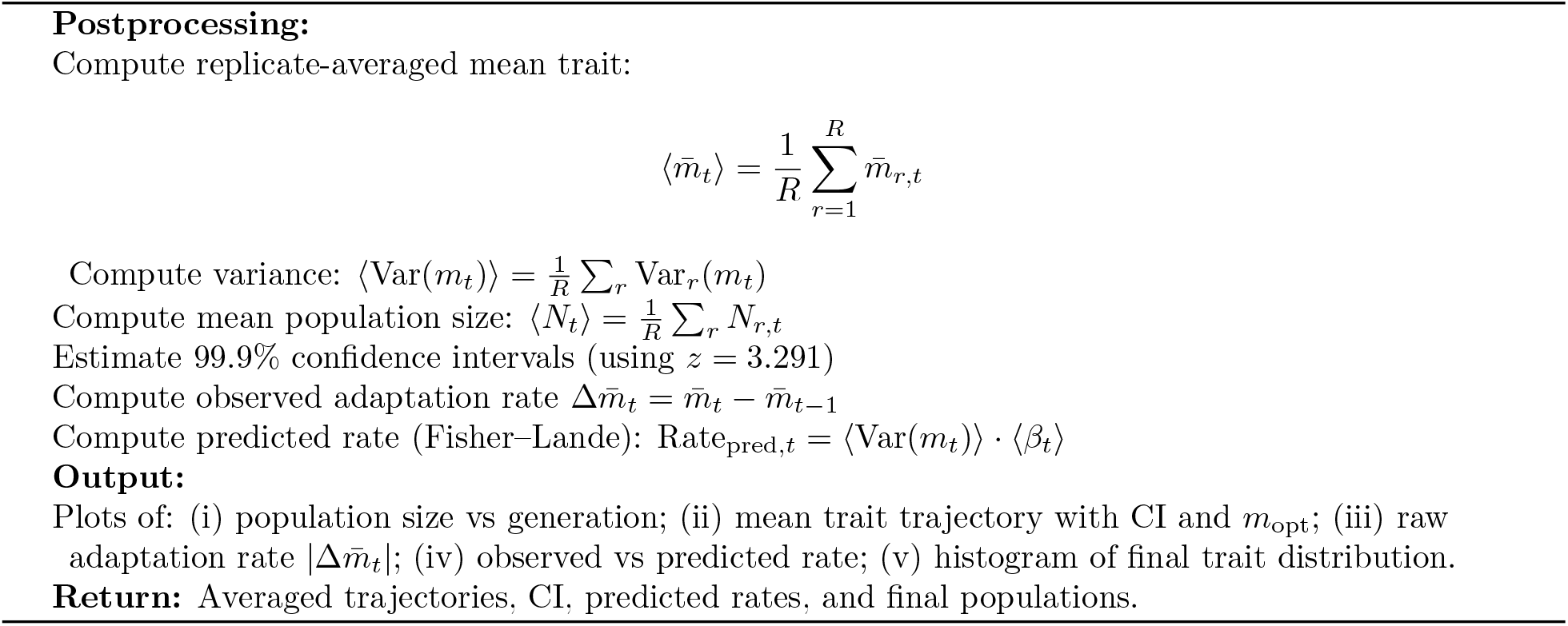

## 3 Quantification of the increased error rate due to perturbation

Values of the optimized driving rate constant and the error rate in the prevailing environment can be calculated using ‘driving function.m’and ‘error calculator.m’:

- Driving rate constant = driving function(f, *K*_*x*_)
- Error rate = error calculator(m, *K*_*x*_)

Increased error rate due to the perturbation can be quantified with following algorithm (sub-script old and new represent the prevailing environment and perturbed environment).

- Optimized driving rate constant (*M*_0,*old*_) = driving function(*f*_*min*_, *K*_*x,old*_)
- Increased error rate (*f*^*′*^) = error calculator(*M*_0,*old*_, *K*_*x,new*_)
- 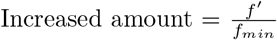

The MATLAB functions ‘driving_function.m’ and ‘error_calculator.m’, which are used to compute the increment in the error rate, are available at: https://github.com/Sashik05/kinetic-proofreading-mutation-robustness.git.

## 4 Enzyme evolution considering identical phenotypic change for entire population and individual phenotypic change

Evolutionary trajectory of *m*_*avg*_ of the population, by assuming that the change in driving rate constant (phenotypic change) in a generation to be uniform across the whole population is illustrated in Fig. S1a.

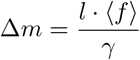

Similarly, Fig. S1b demonstrates the *m*_*avg*_ evolution across generations, when the phenotypic change (driving rate constant) is different for each individual depending on their individual replication error rate.

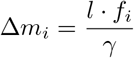

**Supplementary Figure S1:**
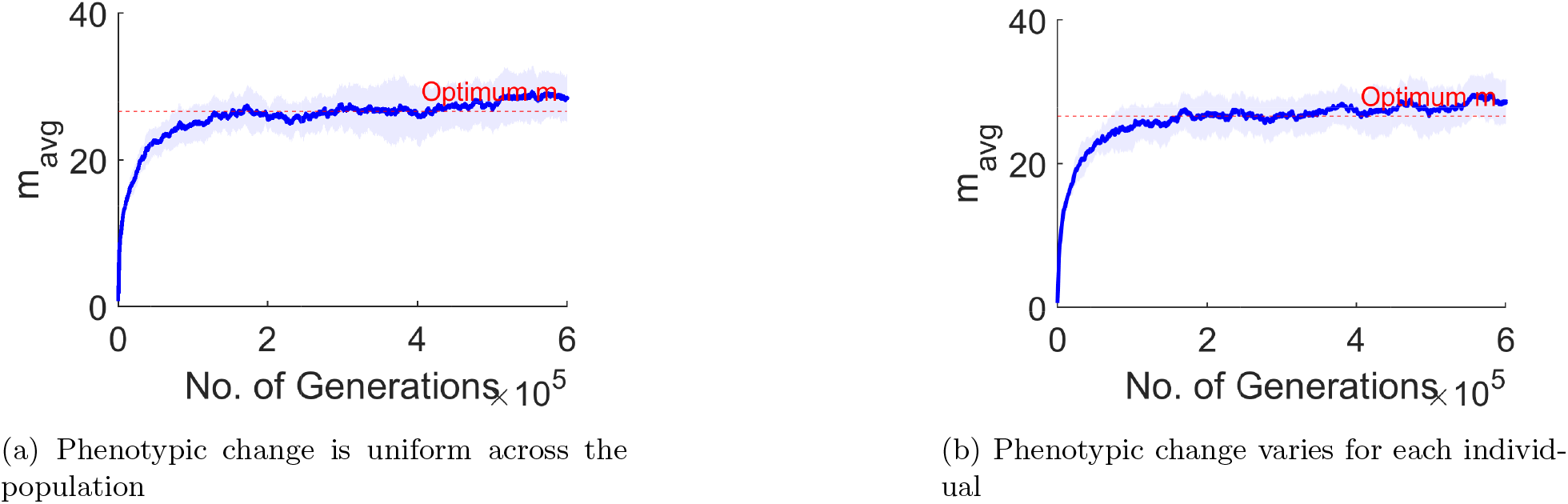
This figure illustrates the adaptive dynamics of the driving rate constant (evolving phenotype) in the perturbed environment. (a) Phenotypic change due to genomic variation is assumed to be uniform across the population. (b) Phenotypic change is different for each individual in a population across generations. However, the adaptive dynamics in both cases are similar.

## 5 Comparison of standard adaptation rate with the observed adaptation rate

To quantitatively evaluate the consistency of our model, we define the following adaptation metrics. Here, we define the effective fitness per generation (*W*) of the evolving trait m. Effective fitness integrates the population’s survival probability with its fecundity, thereby capturing both persistence and reproductive output. Operationally, *W* represents the expected number of individuals, including newly mutated variants, those contribute to the next generation. This measure thus provides a consistent quantitative basis for tracking the course of adaptation. This formulation follows established models of evolutionary prediction (Fisher’s fundamental theorem and Lande’s quantitative genetic framework [1, 2]).

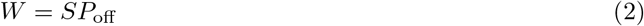

Here, *S* and *P*_off_ denote the mean selection coefficient and the mean offspring probability of the population in a given generation, respectively.

To further quantify evolutionary dynamics and to track the propagation of effective fitness across generations, we define two measures of adaptation rates: the standard and the observed. The standard adaptation rate follows the Fisher/Lande framework, where the expected change in the mean trait 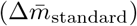 is approximated by the product of trait variance, (Var(*m*)), and the selection gradient, *β*, which measures how fitness changes with the trait [1, 2, 3].

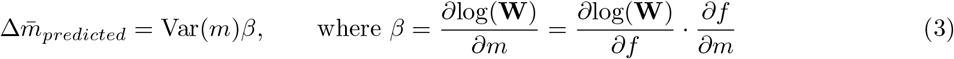

Conversely, the observed adaptation rate 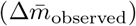 reflects the change in the population mean trait across successive generations:

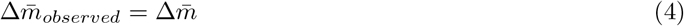

Here, Var(*m*) denotes the trait variance, and 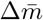 represents the generational change in the population mean trait value.

**Supplementary Figure S2:**
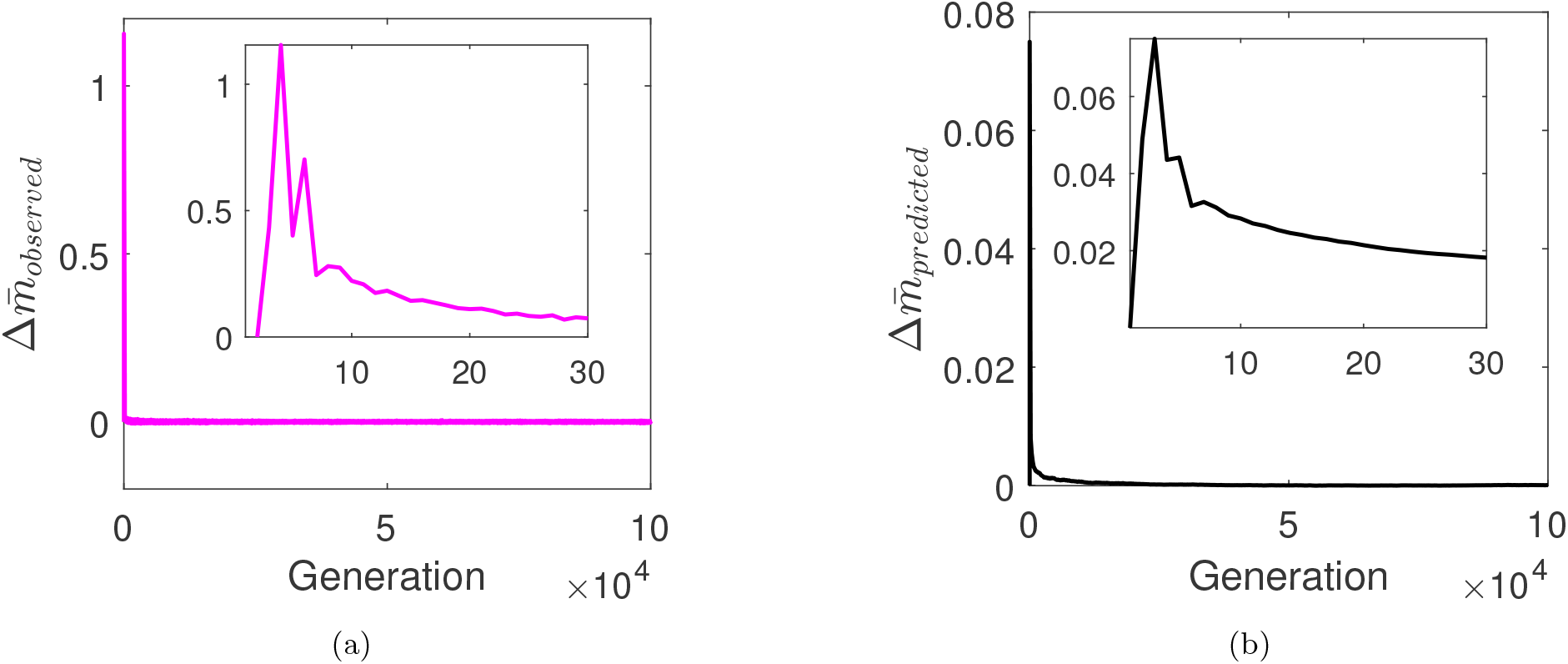
Evolution of the adaptation rate across generations. (a) The observed adaptation rate (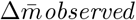 observed, magenta curve) and (b) the predicted adaptation rate (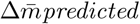 predicted, black curve) exhibit closely aligned trajectories, validating the consistency of the model. In both cases, the adaptation rate rises sharply in the early generations, reflecting the sudden increase in replication error following the environmental (temperature) shift. As the population mean trait approaches the new optimum, the adaptation rate progressively declines, driven by the reduced mutation rate at steady state.

As shown in Fig. S2, both adaptation rate measures follow the same evolutionary trend: an initial sharp rise reflecting the sudden error increase imposed by the temperature shift, followed by a gradual decline as mutations become less frequent and the population mean converges toward the new optimum (*m*_0,new_). This consistent pattern in adaptation rates validates the assumptions of our model.

## 6 Adaptation Dynamics Following Temperature Decline

Following the algorithm in ‘*Evolutionary Framework* ‘, we introduced a perturbation of a temperature shift of Δ*T* = 42 *K* from 335 K (*m*_0,*old*_ = 26.5 *s*^*−*1^, *K*_*x*_ = 42 *s*^*−*1^) to 293 K (*m*_0,*new*_ = 0.6 *s*^*−*1^, *K*_*x*_ = 1 *s*^*−*1^) on a population with initial size 10^3^ and coding region size as 10^6^ bp.

**Supplementary Figure S3:**
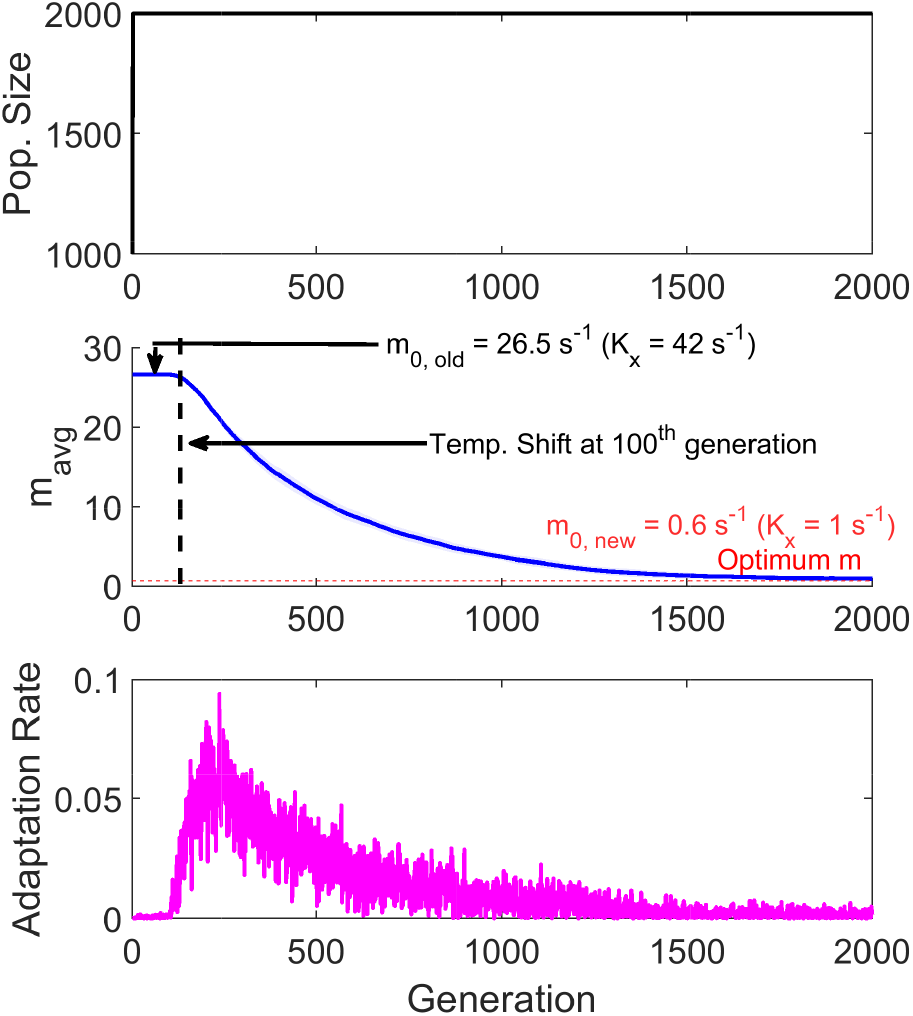
The figure illustrates the adaptation dynamics of a population with a coding-region size of 10^6^ bp following a temperature shift of ΔT = 42 K, from 335 K (m_0,old_ = 26.5 s^−1^, K_x_ = 42 s^−1^) to 293 K (m_0,new_ = 0.6 s^−1^, K_x_ = 1 s^−1^). The black curve shows population growth, the blue curve tracks adaptation toward the new optimal driving-rate constant, and the magenta curve depicts the adaptation rate across generations, representing the change in the phenotypic driving-rate constant per generation.

## 7 Choice of Selection Co-efficient

To associate replication error rates with survival probability, the individual elimination probability is defined as

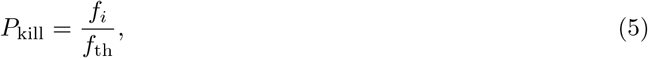

where *f*_*i*_ is the individual error rate (Eq. 3) and *f*_*th*_ = 10^6^ is the threshold beyond which survival becomes unlikely.

The probability of generating viable offspring is expressed as

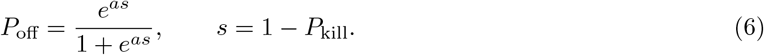

Where ‘s’ is the selection coefficient and defined as the survival probability, while ‘a’ dictates the steepness of the offspring probability curve, discriminating between low and high-fitness individuals.

**Supplementary Figure S4:**
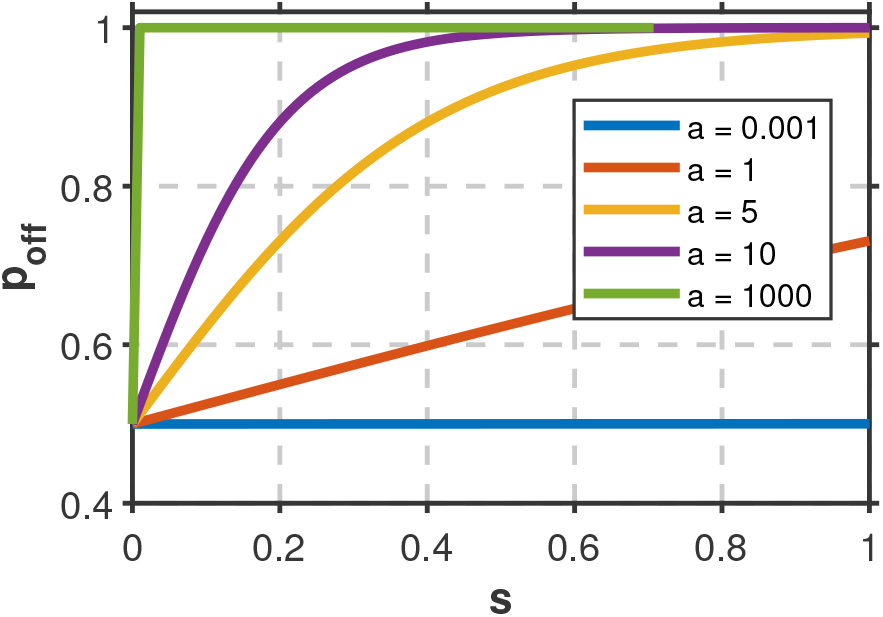
The figure illustrates how the offspring probability (P_off_) varies with the selection coefficient (s) for different values of the steepness parameter a (as defined in Eq. 6). When a is very small or very large (blue and green curves), P_off_ loses its ability to discriminate between high- and low-fitness individuals. Intermediate values of a, particularly around a = 5, provide the strongest discrimination and therefore yield a more biologically meaningful mapping between fitness and reproductive probability.

Fig. S4 illustrates how the steepness parameter *a* shapes the relationship between offspring probability (*P*_off_) and the selection coefficient (*s*). When *a* is very small (e.g., *a* = 0.001), *P*_off_ = 0.5 for all values of *s*, indicating a complete loss of discriminatory power between low- and high-fitness individuals. Conversely, when *a* is very large, *P*_off_ saturates at 1 across the full range of *s*, again eliminating meaningful selection. In contrast, intermediate values of the steepness parameter produce a realistic fitness-to-offspring mapping. As shown in Fig. S4, *a* = 5 yields desirable behavior: as *s →* 0, *P*_off_ *→* 0.5, and as *s →* 1, *P*_off_ *→* 1. Similar qualitative behavior holds for values of *a* near 5 (e.g., *a* = 4, 6, 7), all of which provide adequate discrimination between individuals of differing fitness.

## 8 Dependence of dissociation rate constant (K_x_) with temperature shift

The relationship between dissociation rate constant (*K*_*x*_) with temperature follows the transition state equation:

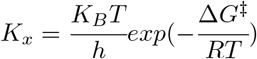

Where Δ*G*^*‡*^ is the free energy of activation, *K*_*B*_ is Boltzmann’s constant (*K*_*B*_ = 1.38 × 10^*−*23^ *JK*^*−*1^), *T* is the absolute temperature, *R* is the universal gas constant (*R* = 8.314 *Jmol*^*−*1^*K*^*−*1^) and *h* is Planck’s constant (*h* = 6.626 × 10^*−*34^ *Js*). Implementing these values and *K*_*x*_ = 2*s*^*−*1^ at 293 *K* from [4, 5], we estimate Δ*G*^*‡*^ = 70.023 *kJmol*^*−*1^.

**Supplementary Figure S5:**
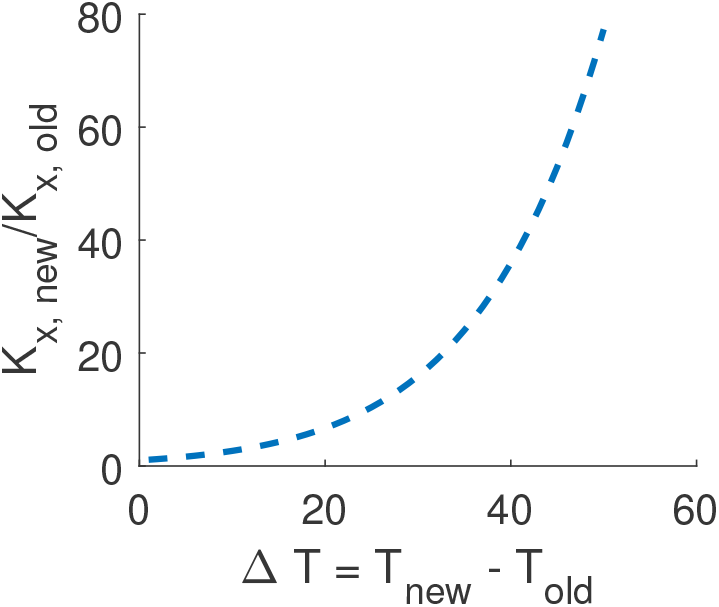
The figure illustrates how the dissociation rate constant varies with temperature. The x-axis denotes the change in temperature (ΔT), and the y-axis shows the corresponding fold-change in the dissociation rate constant.

## 9 Dependence of increased error rate with temperature shift

**Supplementary Figure S6:**
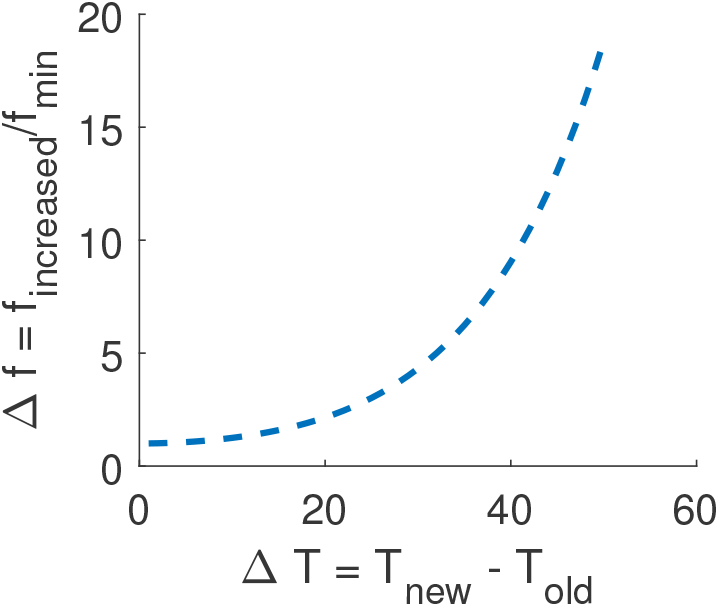
The figure illustrates the increase in replication error rate as a function of temperature shift. The x-axis denotes the magnitude of the temperature change (ΔT), and the y-axis shows the corresponding increase in error rate.

## 10 Evolution under a temperature shift of 10K

**Supplementary Figure S7:**
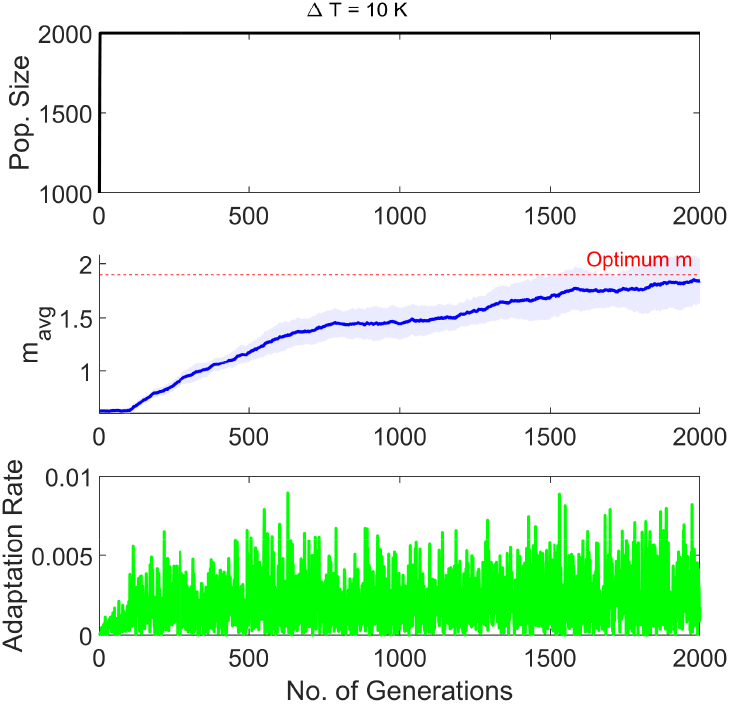
The figure illustrates the evolutionary trajectory of enzyme population under a temperature shift of 10 K.

## 11 Evolution for different P_off_ boundaries

**Supplementary Figure S8:**
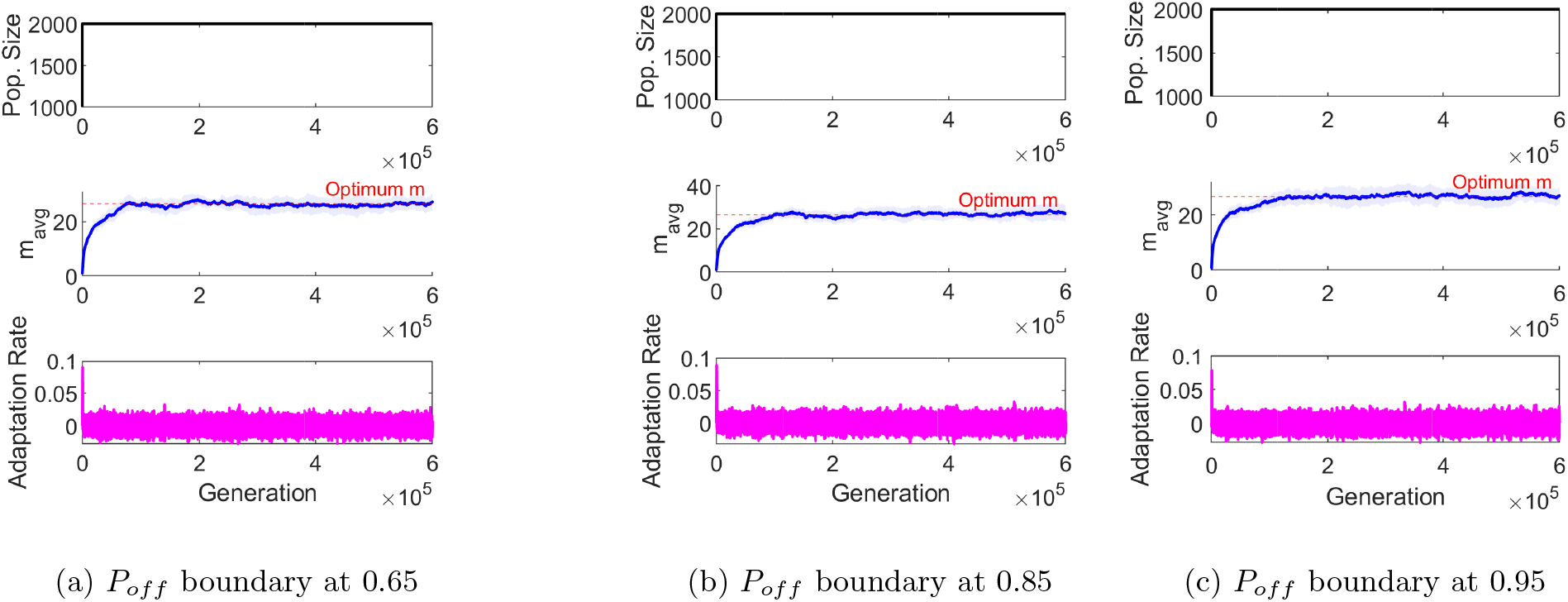
The figure illustrates the adaptive dynamics of a population with an initial size of 10^3^ subjected to an environmental perturbation corresponding to a 42 K temperature shift, evaluated under three different stochastic offspring-probability thresholds (P_off_). Although these boundaries influence the rate of adaptation, with higher thresholds generating more offspring and thereby enhancing selective filtering, they do not alter the qualitative outcome of the evolutionary trajectory. In (a) individuals produce two offspring when P_off_ ≥ 0.65, otherwise one; in (b), the threshold is P_off_ ≥ 0.85; and in (c), P_off_ ≥ 0.95 triggers two-offspring production. These choices introduce stochasticity without changing the overall adaptive response.

## 12 Adaptability of different populations at different temperatures

**Supplementary Figure S9:**
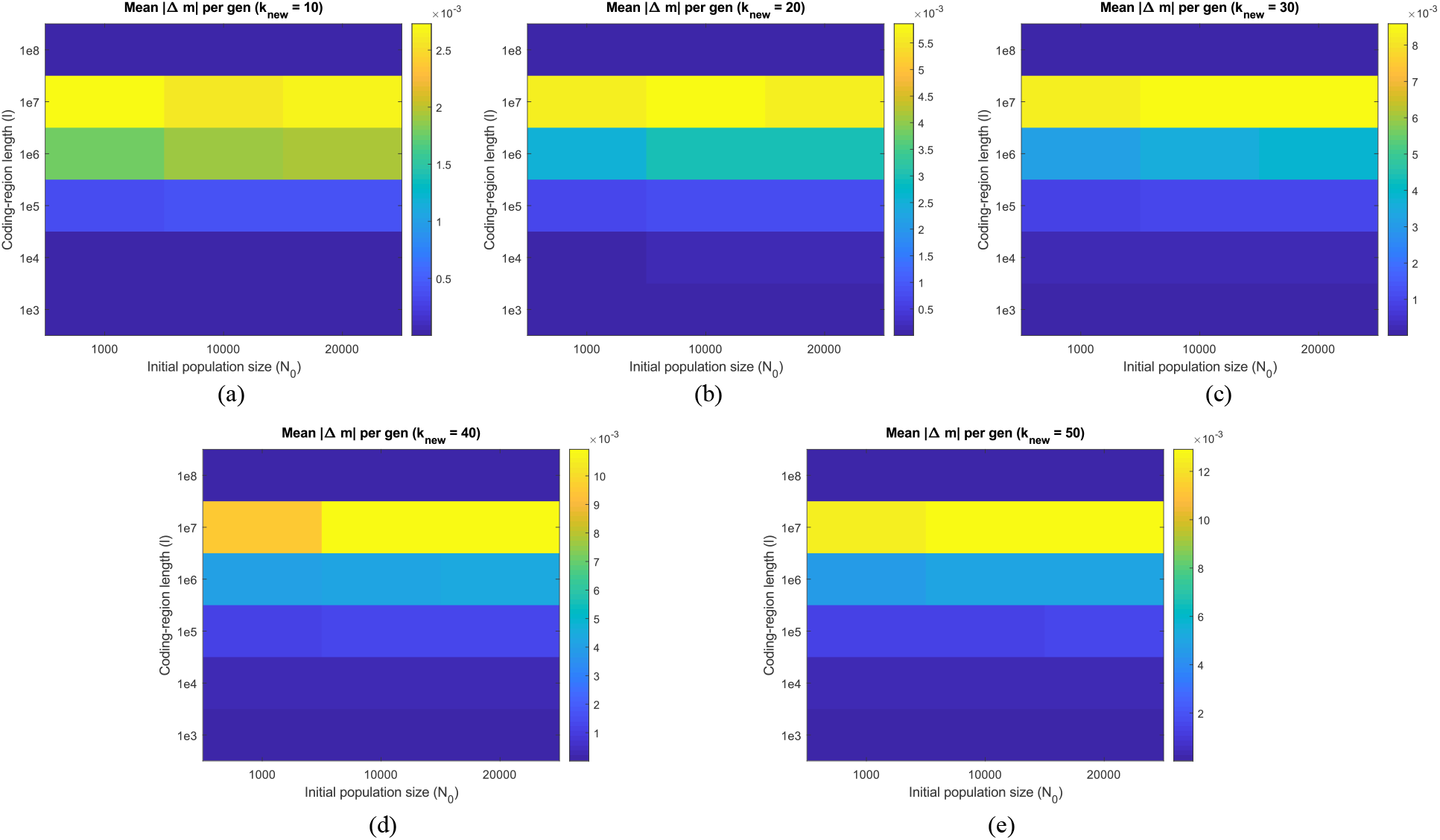
Heatmaps showing the adaptability of populations across combinations of initial population size and coding-region length under different magnitudes of environmental perturbation. Each panel displays the average per-generation change in the phenotypic driving rate constant (adaptability) for three initial population sizes (10^3^, 10^4^, 2 × 10^4^) evaluated across coding-region lengths 10^3^, 10^4^, 10^5^, 10^6^, 10^7^, and 10^8^ bp. Panels correspond to increasing perturbation strength, parameterized by the perturbed dissociation rate constant K_x,new_: **(a)** K_x,new_ = 10 at ΔT = 24.5 K; **(b)** K_x,new_ = 20 at ΔT = 32.7 K; **(c)** K_x,new_ = 30 at ΔT = 37.7 K; **(d)** K_x,new_ = 40 at ΔT = 41.37 K; **(e)** K_x,new_ = 50 at ΔT = 44.27 K. The color bar indicates the mean adaptation rate, quantified as the average change in the driving rate constant per generation. Across all perturbation strengths, populations with coding-region lengths 10^6^ and 10^7^ bp consistently exhibit the highest adaptability.

## 13 Variation of error rate with driving rate constant with discrimination in the association rates

**Supplementary Figure S10:**
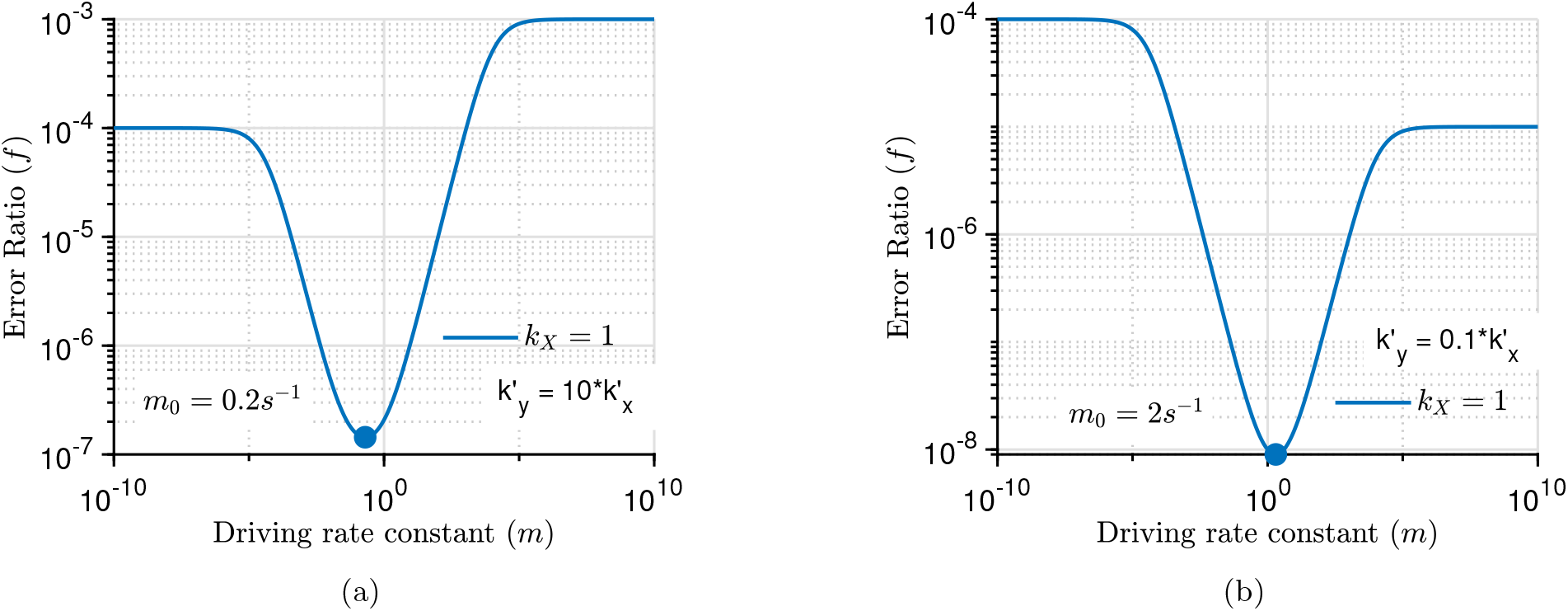
The figure illustrates the variation of the error rate f with the driving rate constant m for a modest difference between the association rates (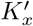 and 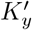) governing the attachment of a new dNTP to the polymerase active site. (a) f vs. m for 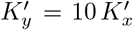 with 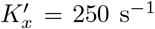; the optimal driving rate constant m_o_ shifts from 0.6 s^−1^ to 0.2 s^−1^. (b) f vs. m for 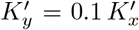 with 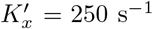; the optimal driving rate constant m_o_ shifts from 0.6 s^−1^ to 2 s^−1^. When the association rate of the incorrect substrate (Y) increases, [EY] > [EX], a smaller optimal driving rate constant is required to balance the discriminatory flux between the two dissociation pathways (steps 1 and 3). Conversely, when the association rate of the incorrect substrate is lower than that of the correct one, [EY] < [EX], the optimal driving rate constant increases to balance the discriminatory flux between the competing pathways.

## 14 Correct product formation rate vs driving rate constant

**Supplementary Figure S11:**
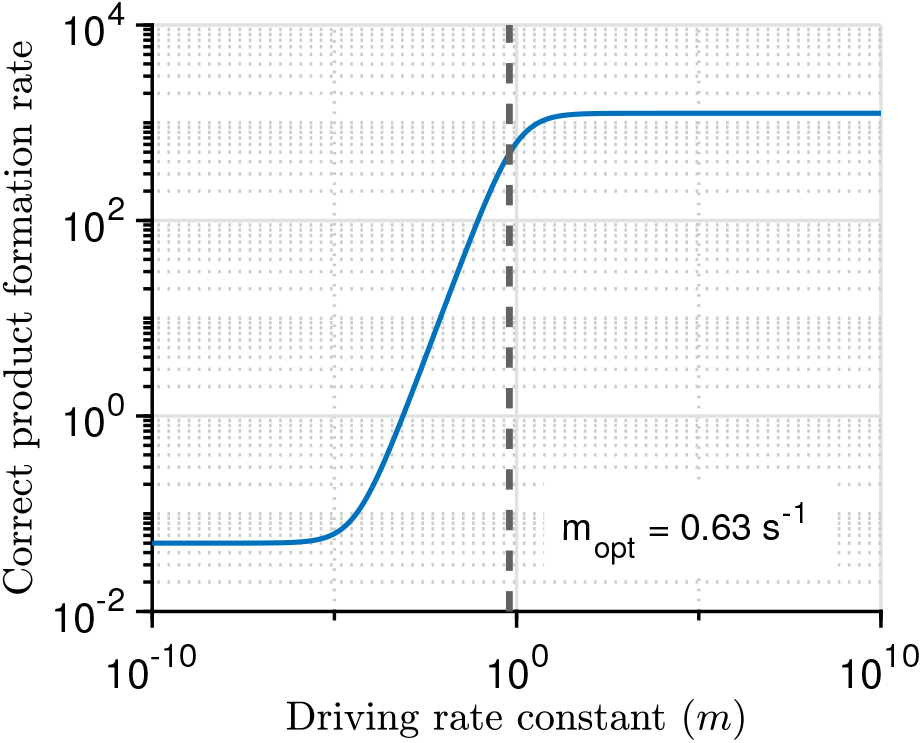
The figure illustrates the variation of the correct product formation rate with the driving rate constant. For very small values of the driving rate constant, the speed remains constant. With further increase in the driving rate constant, the speed increases and attains saturation at a driving rate of 30 s^−1^. The optimal driving rate constant lies near the saturation value of the speed. Here, 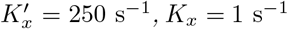, L_x_ = 0.2 s^−1^, and 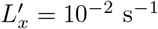.

## 15 Evolution under biased culling

The above evolutionary dynamics is modeled by implementing uniform culling without bias to maintain a finite population size. Under this neutral regulation, the final population exhibited a broader distribution of driving rate constants than the initial population (Fig. 3b), and the mean value *m*_*avg*_ oscillated around *m*_0,new_ after reaching the adapted state in the perturbed environment (Fig. 3c). However, when biased culling was introduced, with viability as the primary and speed (correct product formation rate) as the secondary criterion, *m*_*avg*_ converged to a new value higher than *m*_0,new_ (Fig. S12). Together with Fig. S11, this indicates that the evolved population favors higher speed while remaining within a viable error-rate range, i.e., near the minimum-error regime.

**Supplementary Figure S12:**
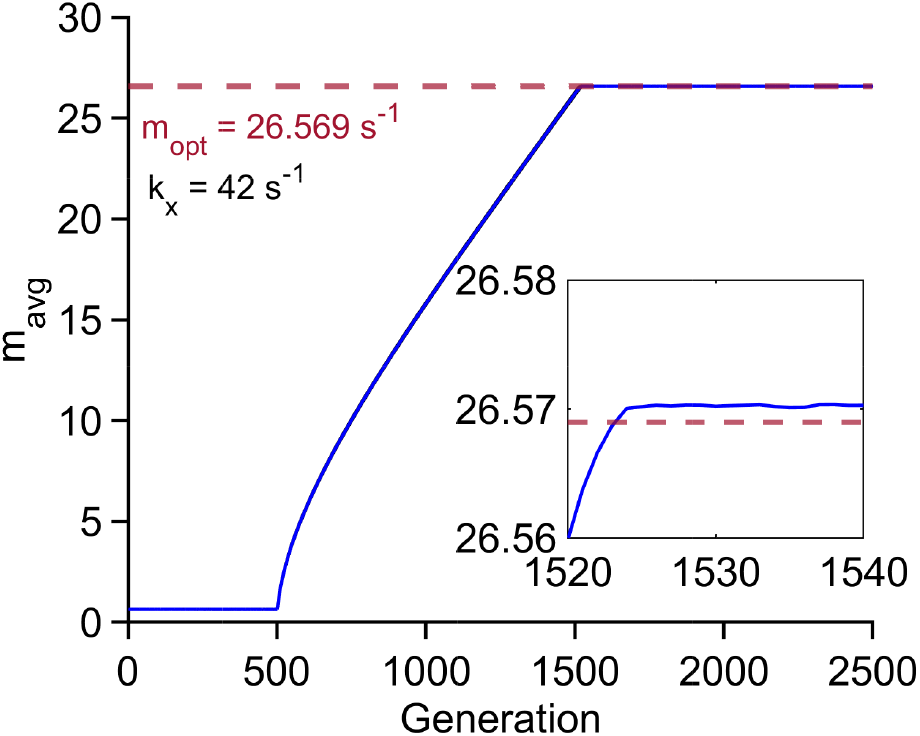
The figure illustrates the evolution of m_avg_ of the population following a perturbation from K_x_ = 1 s^−1^ to K_x_ = 42 s^−1^. A biased culling scheme is used to maintain a finite population size, where viability is given primary priority and the speed (correct product formation rate) secondary priority. We observe that m_avg_ reaches the adaptation state faster than under random culling and maintains an offset toward higher m values, indicating that the population favors speed while preserving viability.

### Kinetic Parameters

**Supplementary Table S1:** Representative kinetic parameters used in the simulations.

| Parameter | Value | Unit |
| --- | --- | --- |
| $K'_X = K'_Y$ | 250 | $\text{s}^{-1}$ |
| $L'_X = L'_Y$ | $10^{-2}$ | $\text{s}^{-1}$ |
| $L_X$ | 0.2 | $\text{s}^{-1}$ |
| $K_X$ | 2 | $\text{s}^{-1}$ |
| $f_0 = \frac{K_X}{K_Y}$ | $10^{-4}$ | – |
| Energetic discrimination factor $\alpha$ | $10^{-4}$ | – |

## 16 Emergent Signatures of Punctuated Evolution

In the molecular evolution of enzymes involved in genomic processes such as DNA replication and protein synthesis, environmental stress leads to an increase in errors within these processes. Increased errors in one genomic process can propagate and amplify errors in others. For example, increased errors in DNA replication intensify the errors in protein synthesis, which is inherently error-prone due to its multistage and complex nature [6]. Consequently, the accumulation of such errors across genomic processes increases the mutational frequency at the organism level, causing the rapid evolution of organisms. Notably, such high genetic variations also often accompany speciation [7].

As enzymes adapt to reduce these errors, the overall mutational frequency stabilizes, allowing the organism to adjust to the perturbed environment and enter a phase of evolutionary stasis. This stasis is maintained until another abrupt environmental perturbation, such as a sudden temperature fluctuation, occurs. During such episodes, the elevated error rates in genomic processes drive rapid bursts of evolution. Thus, the interplay between rapid evolutionary change under elevated mutational frequencies and subsequent stabilization at minimal error rates through readjustment of optimal driving rate constant (see Fig. 3 and 4) reflects the signatures of punctuated equilibrium, as illustrated in Fig. S13.

**Supplementary Figure S13:**
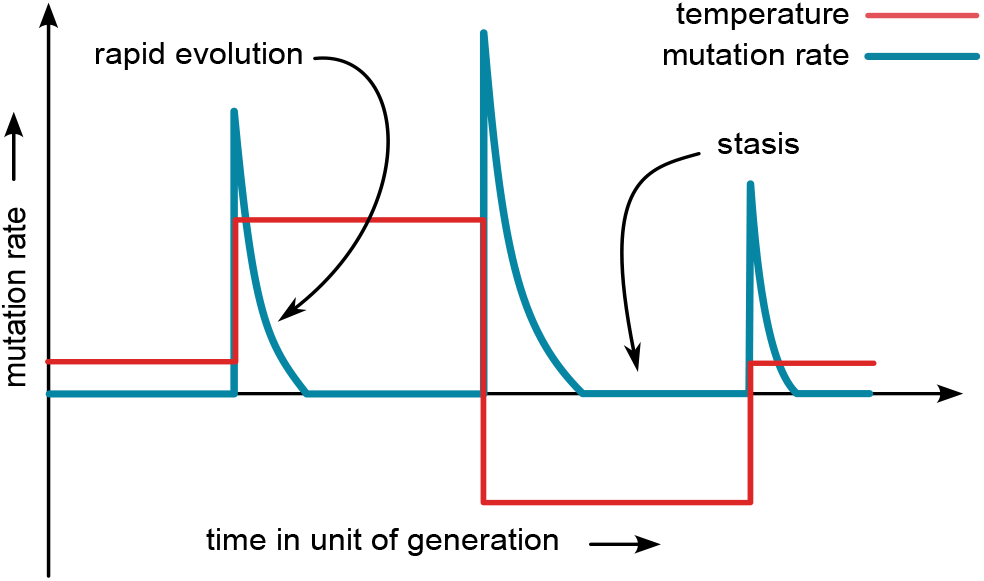
Signature of punctuated equilibrium: Prolonged periods of evolutionary stasis interrupted by rapid bursts of evolutionary change due to sudden environmental perturbations. Temperature is considered here as a proxy for environmental variation, with the red line denoting the temperature changes in the environment over evolutionary timescales. Each abrupt change in temperature increases enzymatic error rates, elevating mutational frequency and driving rapid adaptation towards minimal-error enzymatic configurations. These transitions, shown by the blue curve, can result in the emergence of new traits or even speciation.

Fig. (S13) illustrates that environmental stress, such as temperature fluctuations (red line), disrupts the periods of stasis, leading to rapid evolutionary changes in organisms. The temperature change causes an increase in errors in the genomic processes, which in turn elevates the mutation rate (blue line). This surge in mutations catalyzes swift evolutionary adaptations, as the organism undergoes rapid change to cope with the perturbed environment. Once adaptation minimizes error rates, mutation rate stabilizes, and the system returns to stasis until the next environmental shift. Importantly, this dynamic mirrors the adaptation-rate framework introduced earlier: rapid adaptation corresponds to the steep rise in observed and predicted rates, whereas stasis reflects the gradual decline as the population mean trait approaches the new optimum (Fig. 4).

